# The CRISPR-Cas System differentially regulates surface-attached and pellicle-biofilm in *Salmonella enterica* serovar Typhimurium

**DOI:** 10.1101/2022.01.20.477050

**Authors:** Nandita Sharma, Ankita Das, Pujitha Raja, Sandhya Amol Marathe

**Affiliations:** Department of Biological Sciences, Birla Institute of Technology and Science (BITS), Pilani. Rajasthan, India

**Author notes:** **Corresponding authors:** Sandhya Amol Marathe., Nandita Sharma.

**Keywords:** *Salmonella*, type I-E CRISPR-Cas system, surface-attached biofilm, pellicle-biofilm

## Abstract

CRISPR-Cas (Clustered Regularly Interspaced Short Palindromic Repeats-CRISPR associated) system has been studied for its role in biofilm regulation and expression of outer membrane proteins in *Salmonella*. We investigated the CRISPR-Cas mediated biofilm regulation in *Salmonella enterica* serovar Typhimurium by deleting CRISPR-Cas components, *ΔcrisprI*, *ΔcrisprII, ΔΔcrisprI crisprII,* and *Δcas op.* We determined that the system positively regulates surface-biofilm while inhibiting pellicle-biofilm. In knockout strains, flagellar (*fliC, flgK*) and curli (*csgA*) genes were repressed, causing reduced surface-biofilm. Conversely, they displayed altered pellicle-biofilm architecture possessing bacterial multilayers and a denser ECM with enhanced cellulose and lesser Curli, ergo weaker pellicle. The intracellular cellulose concentration was less in the knockout strains due to upregulation of *bcsC*, necessary for cellulose export. We hypothesized that exported cellulose integrates into the pellicle. We determined that *crp* is upregulated in the knockout strains, thereby inhibiting the expression of *csgD*, hence *csgA* and *bcsA*. The conflicting upregulation of *bcsC*, the last gene of *bcsABZC* operon, could be independently regulated by the CRISPR-Cas system owing to a partial match between the CRISPR-spacers and *bcsC* gene. The CRP-mediated regulation of the flagellar genes in the knockout strains was probably circumvented through the regulation of Yddx governing the availability of σ^28^ factor that further regulates class3 flagellar genes (*fliC, flgK*). Additionally, the variations in the LPS profile and expression of LPS-related genes (*rfaC, rfbG, rfbI*) in the knockout strains could also contribute to the altered pellicle architecture. Collectively, we establish that the CRISPR-Cas system differentially regulates the formation of surface-attached and pellicle-biofilm.

## Introduction

Clustered <u>R</u>egularly <u>I</u>nterspaced <u>S</u>hort <u>P</u>alindromic <u>R</u>epeats (CRISPR)-<u>C</u>RISPR-<u>as</u>sociated (Cas) system bestows adaptive immunity to bacteria against invading mobile genetic elements (MGE) [1]. It captures proto-spacers from invading MGE's and incorporates them in the CRISPR array with the help of Cas proteins [2]. The system has also been implicated in alternative functions like governing virulence and bacterial physiology [3]. In some bacterial species, including *Salmonella*, selective proto-spacers have been traced to chromosomes, thereby supporting a role for the CRISPR-Cas system in endogenous gene regulation [4] [5]. *Salmonella* possesses a type I-E CRISPR-Cas system comprising two CRISPR arrays, CRISPR-I and CRISPR-II, and one *cas* operon [4]. This system has been demonstrated to regulate biofilm formation in *Salmonella enterica* subspecies *enterica* serovar Enteritidis by regulating the quorum-sensing system [6]. It also regulates the expression of outer membrane proteins in serovar Typhi, thereby impacting the biofilm formation and resistance to bile [7].

*Salmonella* is one among the four leading causes of diarrheal diseases worldwide [8]. Salmonellosis, a disease caused by *Salmonella*, presents a formidable threat to humans while causing typhoid fever in ~14.3 million individuals with 135,000 estimated deaths worldwide[9]. *Salmonella enterica* forms biofilms on medically important surfaces like medical devices (catheters, endoscopy tubes, etc.) and gallstones[10], complicating the treatment processes. Biofilm formation on food surfaces has also been correlated to *Salmonella's* persistence, thereby safeguarding it throughout food processing[11]. Biofilm formation on cholesterol-rich gallstones is conceived as a significant factor influencing the establishment of a chronic carrier state, accounting for 1-6 % of total typhoid cases[12],[13]. Biofilm aids *Salmonella* virulence by facilitating evasion of the hosts' immune response and increasing antibiotic tolerance as biofilms are impenetrable to antibiotics.

In this study, we evaluated if and how the endogenous CRISPR-Cas system plays a role in regulating the biofilm formation of *Salmonella enterica* subspecies *enterica* serovar Typhimurium (*S*. Typhimurium). We found that the CRISPR-Cas system differentially regulated surface-attached and pellicle biofilm formation by altering the expression of biofilm-associated genes.

### Importance

In addition to being implicated in bacterial immunity and genome editing, the CRISPR-Cas system has recently been demonstrated to regulate endogenous gene expression and biofilm formation. While the function of individual *cas* genes in controlling *Salmonella* biofilm has been explored, the regulatory role of CRISPR arrays in biofilm is less studied. Moreover, studies have focused on the effects of the CRISPR-Cas system on surface-associated biofilms, and comprehensive studies on the impact of the system on pellicle biofilm remain an unexplored niche. We demonstrate that the CRISPR array and *cas* genes modulate the expression of various biofilm genes in *Salmonella*, whereby surface- and pellicle-biofilm formation is distinctively regulated.

## Material and Methods

### Bacterial strains and culture conditions

*S.* Typhimurium str. 14028s was used as a parent strain (wildtype, WT 14028s). The wildtype, CRISPR, and *cas* knockout strains (the knockout construction is explained below) and their corresponding complement strains were routinely grown in Luria-Bertani (LB, HiMedia) with appropriate antibiotics (Supplementary table 1) at 37°C, 120 rpm. The bacterial strains were also sub-cultured and grown in biofilm-media (LB without NaCl: 1% tryptone, 0.5% yeast extract) for observing growth patterns up to 12 h.

### Construction of CRISPR and *cas* operon knockout strains

We generated the CRISPR and *cas* operon knockout strains, *ΔcrisprI* (CRISPR-I array deleted)*, ΔcrisprII* (CRISPR-II array deleted)*, ΔΔcrisprI crisprII* (CRISPR-I and CRISPR-II arrays deleted), and *Δcas op.* (*cas* operon deleted) using a one-step gene knockout strategy described by Datsenko *et al*.[14]. A phage lambda-derived Red recombination system (supplied on the pKD46 plasmid) was used to replace the desired genes in *S.* Typhimurium str. 14028s with a chloramphenicol resistance cassette. The double knockout strain, *ΔΔcrisprI crisprII,* was constructed by replacing the *crisprI* gene with a kanamycin resistance cassette in the *ΔcrisprII* strain. The primers used for knockout generation are listed in supplementary table 2.

### Generation of complement strains for the knockout

The *crisprI* and *crisprII* genes were amplified using the respective cloning primers listed in supplementary table 2. The amplified products were cloned into *Bam*HI and *Hind*III sites of pQE60 (A kind gift from Prof. Dipshikha Chakravortty, Indian Institute of Science, India). The positive constructs were transformed into the respective knockout strains to obtain corresponding complement strains, *ΔcrisprI* + p*crisprI* and *ΔcrisprII* + p*crisprII*.

### Biofilm quantification using crystal violet (CV) assay

#### Tube biofilm assay

Overnight grown bacterial cultures were subcultured at 1:100 ratios in LB supplemented with 3% Ox Bile (HiMedia). These cultures were added in 1.5 mL microcentrifuge tubes coated with 1 mg cholesterol and subsequently incubated at 37°C under static conditions for 96 h. Every day, the media was replaced with fresh media (LB+3% Ox-bile). The biofilms were quantified using a CV assay.

#### Ring and pellicle biofilm

Overnight grown bacterial cultures were subcultured at 1:100 ratio in LB without NaCl media in test-tube and incubated at 25°C under static conditions for 24 h, 48 h, and 96 h. The biofilms were quantified using a CV assay.

#### Crystal violet (CV) assay

The biofilms formed were given washes with phosphate-buffered saline (PBS), dried at 56°C for 30 mins, and stained with 1% (w/v) CV solution for 20 mins. After washing with distilled water, biofilms were quantified by solubilizing the biofilm-bound CV with 30% (v/v) glacial acetic acid and recording the absorbance of the solution at 570 nm using Multiskan GO (Thermo Scientific, USA).

### Biofilm dry mass and viability assay

Biofilm dry mass was estimated by recording the weight of the pellicle biofilms dried in a hot air oven at 56°C.

#### Resazurin-based viability assay

The pellicle biofilms were washed twice with distilled water and stained with resazurin (HiMedia) dye (0.337 mg/mL) for 30 mins at room temperature (RT). The resazurin fluorescence was measured using Fluoroskan Ascent^®^ (Thermo Scientific, USA) at excitation (λ_Ex_) 550 nm and emission (λ_Em_) of 600 nm.

### Biofilm architecture using field emission scanning electron microscopy (SEM)

The pellicle biofilms were allowed to form in the glass tube containing an immersed glass slide. The pellicle biofilms fixed with 2.5% glutaraldehyde were dehydrated with increasing ethanol concentrations. The samples were air-dried, sputter-coated with gold, and visualized with FEI ApreoS Field Emission Scanning Electron Microscope (Oxford Instruments, Netherland).

### Confocal laser scanning microscopy (CLSM) for pellicle biofilm

The pellicle biofilm was stained with 5 μM SYTO 9 (Thermo Scientific), 5 μM Propidium Iodide (PI) (Thermo Scientific), and 50 μM Calcofluor white (SIGMA-ALDRICH) solution for 30 mins, in the dark at RT. Slides were imaged with an LSM 880 Confocal Microscope (Zeiss, Germany) using Z-stack (ZEN 2.3 SPI).

### Motility Assay

Five microlitres of overnight cultures were spot inoculated at the center of swarm petri-plates (20 g/L Luria Broth, 0.5% (w/v) agar and 0.5% (w/v) glucose). After 45-50 mins of air drying, the plates were incubated at 37°C for 9 h. The swarm rate was estimated by calculating the radius of the growth front using Image J Software (U. S. National Institutes of Health, USA).

### Evaluation of the expression of flagellar proteins

Planktonic bacterial cells and pellicle biofilms were lysed in Laemmli buffer. Pellicle biofilms (96 h) were homogenized with TissueLyser LT (QIAGEN, Germany) at 50 kHz for 10 mins. An equal amount of each lysate (50 μg protein from planktonic and 400 μg from pellicle biofilm) was processed for immunoblotting using an anti-flagellin (DIFCO) antibody. The immunoblots were developed, and images were captured with the ChemiDoc XRS+ system (Bio-Rad Laboratories, USA). Each immunoblot band was normalized to coomassie stained bands, and the relative ratio of each with WT was quantified using Image Lab software (Bio-Rad Laboratories, USA).

### Cellulose Determination

Cellulose dry weight estimation, calcofluor binding, and anthrone assay was used to estimate cellulose content in the pellicle biofilm. For cellulose dry weight estimation, pellicle biofilms were washed twice with distilled water and hydrolyzed with 0.1 M sodium hydroxide (NaOH) at 80°C for 2 h. The samples were dried and weighed.

#### Cellulose quantification by calcofluor

The pellicle biofilms were rinsed twice with distilled water, stained with 50 μM calcofluor white stain (SIGMA-ALDRICH) for 40 mins in the dark at RT. The bound calcofluor was measured at excitation (λ_Ex_) 350 nm and emission (λ_Em_) of 475 nm with VICTOR 3 1420 Multilabel Counter (PerkinElmer, USA).

#### Cellulose quantification by anthrone

The bacterial pellets from planktonic cultures were resuspended in 300 μL of an acetic-nitric reagent and incubated for 30 mins at boiling temperatures. The pellets were then washed twice with sterile water, followed by adding 67% sulphuric acid with intermittent mixings, and incubated at RT for 1 h. The samples were placed on an ice bath, and 1 mL of cold anthrone reagent (FISHER SCIENTIFIC) was added and mixed gently. The tubes were incubated in a boiling water bath for 15 mins, after which they were placed on ice. The absorbance at 620 nm was recorded with Multiskan GO.

### Whole-cell Congo red depletion assay

The planktonic cultures grown for 48 h under static conditions were pelleted at 10,000 x g, 5 mins and resuspended Congo red solution (10 μg/mL). After 10 mins incubation at RT, the cells were centrifuged at 10,000 x g, 10 mins. The absorbance of the supernatant was measured at 500 nm with Multiskan GO.

### Curli estimation by ThT fluorescence

The pellicle biofilms were lysed with a lysis buffer (Tris EDTA, pH 7.5 and 2% SDS) at 95°C for 45 mins. The insoluble pellet was washed twice with autoclaved water and resuspended in PBS containing DNase (1 mg/mL, HiMedia) and RNase (20 mg/mL, HiMedia). After 6 h incubation at RT, the samples were treated with 2 μM of ThT (SIGMA-ALDRICH) for 15-20 mins in the dark. The absorbance was measured at excitation (λ_Ex_) 440 nm and emission (λ_Em_) 482 nm with the VICTOR 3 1420 Multilabel Counter.

### Quantitative real-time (q-RT) PCR

Total RNA from 24 h bacterial culture in LB without NaCl was isolated from bacteria using TRIzol reagent (HiMedia) and cDNA synthesized using ProtoScript^®^ II Reverse Transcriptase (NEB). qRT-PCR was performed using PowerUp™ SYBR™ Green Master Mix (Thermo Fisher Scientific). Relative expression of the gene was calculated using the 2^-ΔΔCt^ method by normalizing to reference gene *rpoD.* The primers used in RT-qPCR are listed in supplementary table 2.

### Statistical analysis

Statistical analysis was performed using Prism 8 software (GraphPad, California). Unpaired Student's *t* test was performed. Error bars indicate SD. Statistical significance: *, P≤ 0.05, **, P≤ 0.01, ***, P≤ 0.001, ****, P<0.0001, ns = not significant.

## Results

### CRISPR-Cas knockout strains show temporal variations in the biofilm formation

We tested the biofilm-forming ability of the CRISPR and *cas operon* knockout strains (*ΔcrisprI, ΔcrisprII, Δcas op.,* and *ΔΔcrisprI crisprII*) of *S*. Typhimurium str. 14028s under gall-stone mimicking conditions. At the end of the 96 h, all the knockout strains showed reduced biofilm formation compared to WT (Fig.1A). The phenotypes exhibited by the knockout strains were restored on the complementation of corresponding genes in *ΔcrisprI* and *ΔcrisprII* (Fig. 1A). This confirms that the gene deletions were clean without any side effects. Next, a time-dependent study determining the biofilm formation by the knockout strains in low osmotic conditions (LB without NaCl) showed temporal variations in biofilm phenotypes compared to that of the WT (Fig.1B). The knockout strains formed a thin biofilm ring on the solid-liquid-air interface (surface biofilm) at 24 h (Fig.1B) and 96 h (Supplementary figure, Fig. S1A).

**Figure 1:**
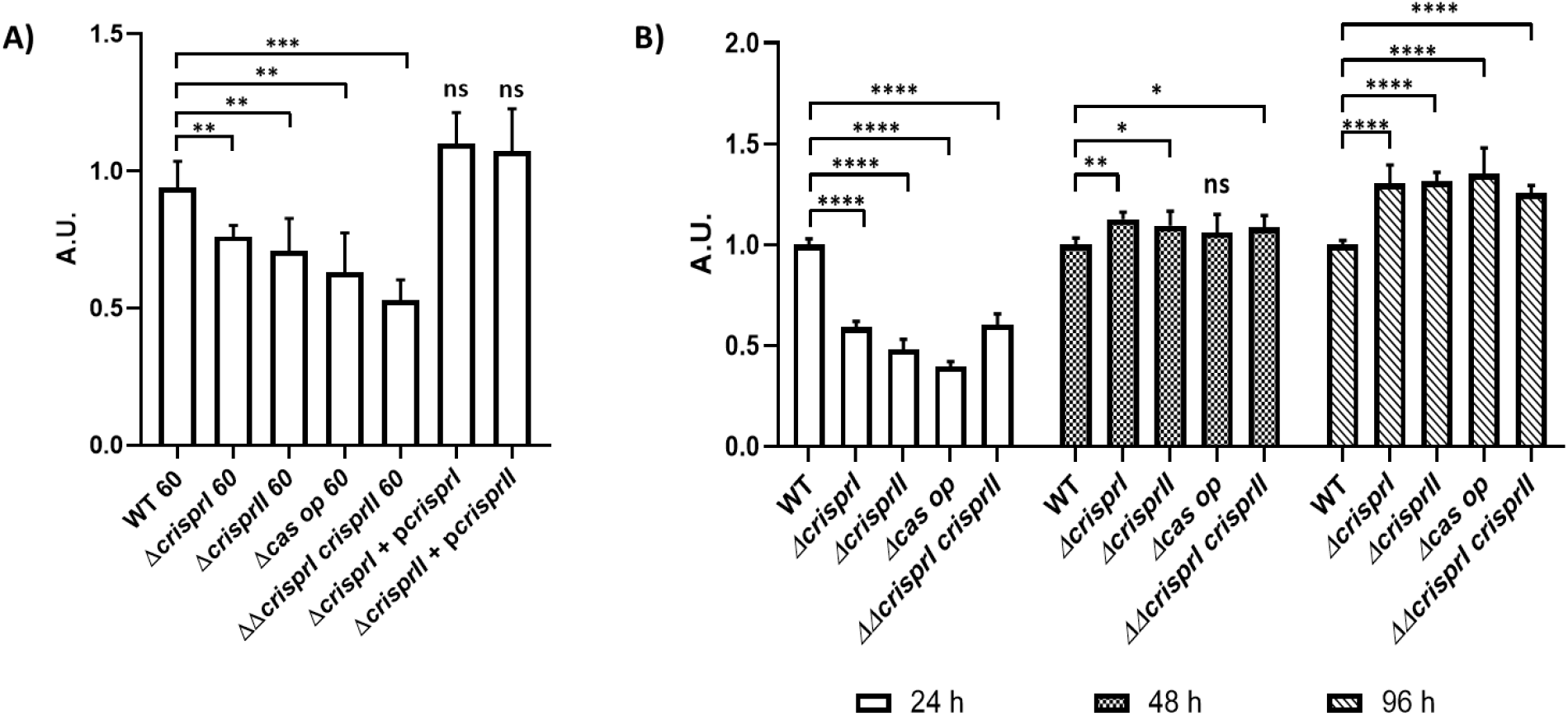
The CRISPR-Cas system knockout strains of *S. enterica* subsp. *enterica* serovar Typhimurium 14028s showed reduced biofilm formation under gallstone mimicking condition (A), while these strains showed temporal variations in the biofilm under low osmotic condition (B). **A.** Wild-type, CRISPR, and *cas operon* knockout strains transformed with empty vector, pQE60 (WT60, *ΔcrisprI 60, ΔcrisprII 60, Δcas op 60,* and *ΔΔcrisprI crisprII 60*), and the complement strains (*ΔcrisprI+*p*crisprI, ΔcrisprII+*p*crisprII*) were cultured in cholesterol coated microcentrifuge tubes in LB media for 96 h at 37°C, static conditions. **B.** The *S.* Typhimurium strain 14028s wildtype (WT), CRISPR (*ΔcrisprI, ΔcrisprII,* and *ΔΔcrisprI crisprII*) and *cas operon (Δcas op*) knockout strains were cultured in LB without NaCl media for different periods (24 h, 48 h, and 96 h) at 25°C, static condition. The biofilm formation in all the cases was estimated using a crystal violet staining assay. The graphs represent OD_570nm_ for each strain, normalized by OD_570nm_ of WT. An unpaired t-test was used to determine significant differences between the WT and knockout strains. Error bars indicate SD. Statistical significance: *≤ 0.05, **≤ 0.01, ***≤ 0.001, ****<0.0001, ns = not significant. A.U., arbitrary units.

However, as time progressed, the knockout strains displayed a gradual increase in biofilm formation, with a significantly high biofilm at 96 h (Fig.1B, and Supplementary figure, Fig. S1B). The difference in observed biofilm phenotype was not accredited to the difference in bacterial growth as testified by the similar growth patterns of all the strains in LB without NaCl media (Supplementary figure, Fig. S2).

### Scanning Electron Microscopy (SEM) depicts the difference in the biofilm architecture of CRISPR-Cas knockout strains

SEM was used to investigate the biofilm architecture at early (24 h) and late (96 h) time points. At 24 h, the micrographs of WT showed more aggregated and tightly packed bacterial cells covering the large surface area (Supplementary figure, Fig. S3). In contrast, the micrographs of all the knockout strains showed patchy bacterial aggregates (Supplementary figure, Fig. S3). Distinct bacterial cells were more evident in *Δcas op*. Small dome-like structures were observed only in the WT micrograph, indicating the formation of the multilayered structure. The biofilm formed by the knockout strains displayed clumped cells without any slimy material in their vicinity. Interestingly, a few elongated cells (marked in micrograph) were observed in the knockout strains at 24 h (Supplementary figure, Fig. S3).

SEM analysis of 96 h pellicle biofilm revealed that, in general, the air-exposed side of the pellicle biofilm had a dry but smooth mat-like structure composed of dense fibrous networks with tightly packed bacterial cells. However, compared to WT biofilm, the biofilms formed by knockout strains had thicker ECM coatings and consisted of 'hilly' structures of different sizes (arrow-heads, Fig.2A). The liquid-submerged side of the pellicle biofilm was rough, consisting of a dome” and valley-like arrangement made up of loosely packed bacterial cells entrapped in EPS. The knockout strains also displayed discrete regions with EPS lumps (marked in micrographs) and pronounced bacterial density (Fig.2B).

**Figure 2:**
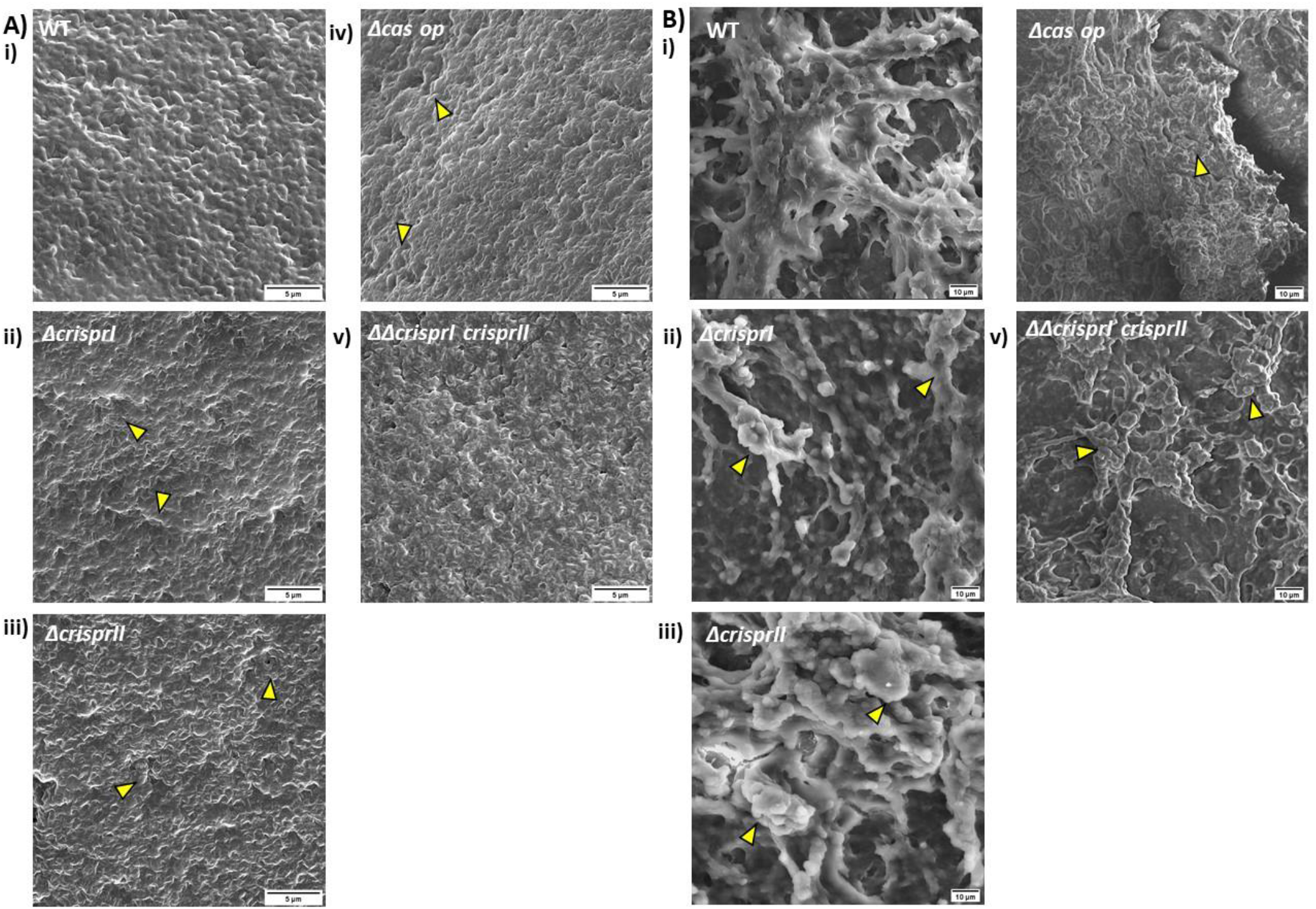
Morphology of air-exposed side (A) and liquid-submerged side (B) of pellicle biofilm at 96 h. The strains were grown in LB without NaCl media for 96 h, at 25°C, static conditions. The pellicle biofilms formed, fixed using 2.5% glutaraldehyde, were dehydrated with increasing ethanol concentrations. SEM image analysis depicts the difference in the pellicle biofilm architecture of CRISPR-Cas knockout (*ΔcrisprI, ΔcrisprII, Δcas op., and ΔΔcrisprI crisprII*) strains, and that of the wildtype (WT), for both air-exposed side (captured at 10,000x magnification), and liquid-submerged (captured at 2500x magnification) side of pellicle biofilm. The air-exposed surface of the pellicle biofilm of CRISPR-Cas knockout strains had denser mat-like ECM. It consisted of "hilly" structures (marked with arrow-heads), indicating more layering of the biofilm. The liquid-submerged surface of the pellicle biofilm of CRISPR-Cas knockout strains had more EPS lumps (marked with arrow-heads) than wildtype. Images were scaled to bar.

### Factors contributing to differential biofilm formation by CRISPR-Cas knockout strains

To understand the knockout strains' temporal variations in biofilm formation, we assessed the expression of essential biofilm components like flagella, cellulose, LPS, and curli.

#### CRISPR-Cas knockout strains show reduced motility and flagellin expression

Motility is crucial for forming surface-associated multicellular communities by several bacteria, including *Salmonella*. It helps in the initial surface colonization during biofilm formation[16]. As the CRISPR and *cas* deletion mutants showed reduced biofilm formation at 24 h (early time-point), we assessed their motility using a swarming assay. There was at least 20% reduction in swarming rates of all the knockout strains compared to WT (Supplementary figure, Fig. S4 and Fig. 3A). The complementation of *ΔcrisprI* and *ΔcrisprII* with corresponding genes restored the defect in their motility (Supplementary figure, Fig. S4, and Fig. 3A). We next analyzed the expression of flagellin protein (FliC) for the planktonic and pellicle bacteria. The immunoblot analysis revealed that the FliC expression in planktonic bacteria was less for knockout strains than that of WT. However, in the 96 h pellicle, no FliC expression was observed in all the strains (Fig.3B).

**Figure 3:**
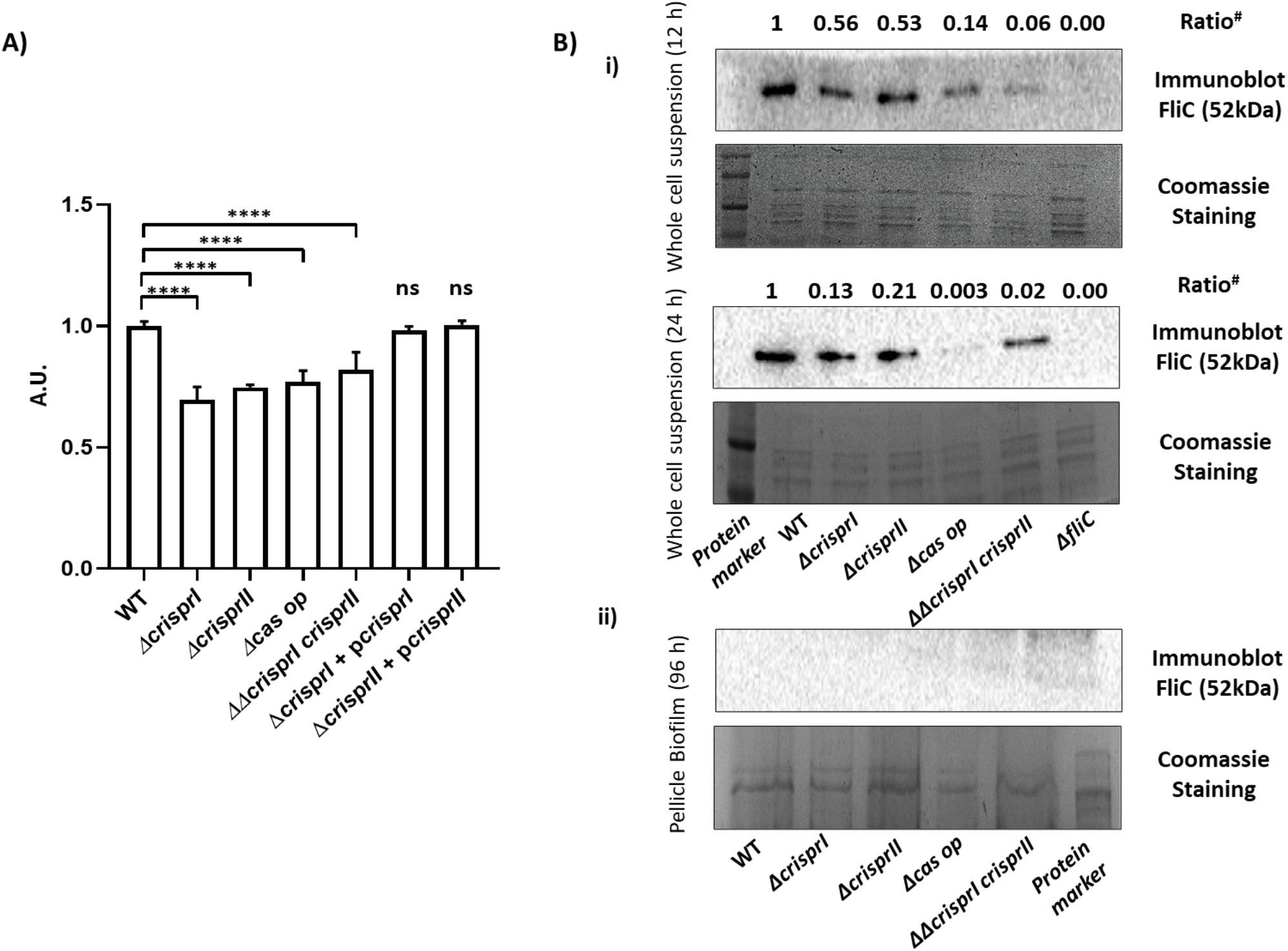
Reduced swarming motility (A), and expression of flagellar protein, FliC (B) was observed in the CRISPR-Cas system knockout strains. **A.** Overnight cultures were point inoculated on swarm agar plates and incubated at 37°C for 9 h. Swarming rate (cm/h) of the wildtype (WT), the knockout strains (*ΔcrisprI, ΔcrisprII, Δcas op,* and *ΔΔcrisprI crisprII*), and the complement strains (*ΔcrisprI* + p*crisprI, ΔcrisprII*+ p*crisprII*) was calculated. The graph represents the swarming rate (cm/h) relative to that of WT. **B.** The strains were grown in LB without NaCl media for different periods (12 h, 24 h, and 96 h), at 25°C, static conditions. The expression of the flagellar protein in planktonic bacteria (B(i)) at early time points (12 h and 24 h), and in pellicle biofilm (B(ii)) at a late time point (96 h) was assessed using western blot analysis with antibodies against FliC. Even at higher protein concentration, FliC was not detected in the blot for pellicle sample of any strain, indicating repression of FliC expression in the pellicle. *ΔfliC* was used as a negative control. An unpaired t-test was used to determine significant differences between the WT and knockout strains. Error bar indicates SD. Statistical significance: *≤ 0.05, **≤ 0.01, ***≤ 0.001, ****<0.0001, ns = not significant. A.U., arbitrary units. # ratio: 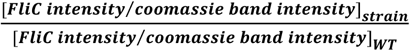

#### Deletion of CRISPR-Cas genes affects the LPS structure

The reduction in swarming motility in the knockout strains is not consistent with FliC expression. For example, expression of FliC protein was minimum in the *ΔΔcrisprI crisprII* strain, but its swarming rate was not the lowest. This anomaly could partially be attributed to the variations in the wettability factor, like LPS that governs the swarming rate. Additionally, the O-antigen of LPS plays a crucial role in biofilm formation [17], and Gram-negative bacteria modify their LPS while in the biofilm [18]. Thus, we assayed the LPS profile of all the knockout strains and compared it with that of the WT (Supplementary figure, Fig. S5). The intensity of the lipid-A band was similar in all the strains, except for *Δcas op.* and *ΔΔcrisprI crisprII*. O-antigen profile showed variations, where the ladder-like banding patterns in *ΔcrisprII* and *ΔΔcrisprI crisprII* were of less intensity than other strains. The band corresponding to very long O-antigen was absent in *ΔcrisprI*, whereas the WT and *Δcas op.* bands had comparable intensities. The very long O-antigen band intensity was similar for *ΔΔcrisprII* and *ΔΔcrisprI crisprII* but was less than that of WT. As for the banding pattern of core glycoforms, *ΔΔcrisprII* and WT were similar to *ΔΔcrisprI crisprII* and *Δcas op.,* respectively. *ΔcrisprI* had a distinct pattern of core glycoforms.

All these observations point to alterations of the O-antigen chain in the knockout strains during biofilm formation.

#### The CRISPR-Cas knockout strains show increased pellicle formation due to increased bacterial biomass and its respective components

The dry weights of the pellicle biofilms by all the knockout strains were similar to that of the WT at 48 h, whereas it was significantly higher at 96 h (Fig. 4A). The temporal variations in the dry weight of all the strains were similar to that of the biofilm formation as estimated using crystal-violet assay. As the dry mass comprises bacterial cells and ECM, we independently assessed the bacterial cell mass (by assessing viability) and concentration of the ECM components. The resazurin cell viability assay results show that the knockout strains are more viable than WT (Supplementary figure, Fig. S6A), hinting at more bacterial mass. We also validated high bacterial abundance within biofilms of knockout strains using SYTO9 staining. Biofilms of all the knockout strains had higher SYTO9 intensity than the WT (Supplementary figure, Fig. S6B) suggesting higher bacterial numbers. Further, the SYTO9: PI ratio was more in the pellicles of the knockout strains, except *Δcas op.,* indicating the presence of more viable bacteria (Fig.4B and Supplementary figure, Fig. S7B). As per the CLSM Z-stack images of the pellicle biofilm, the knockout strains had fewer dead cells near the air-exposed surface than WT (Supplementary figure, Fig S7A). The pellicle biofilm thickness observed by CLSM were 82 μm, 96 μm, 88 μm, 112 μm and 124 μm for WT, *ΔcrisprI, ΔcrisprII*, *Δcas op. and ΔΔcrisprI crisprII* respectively.

**Figure 4:**
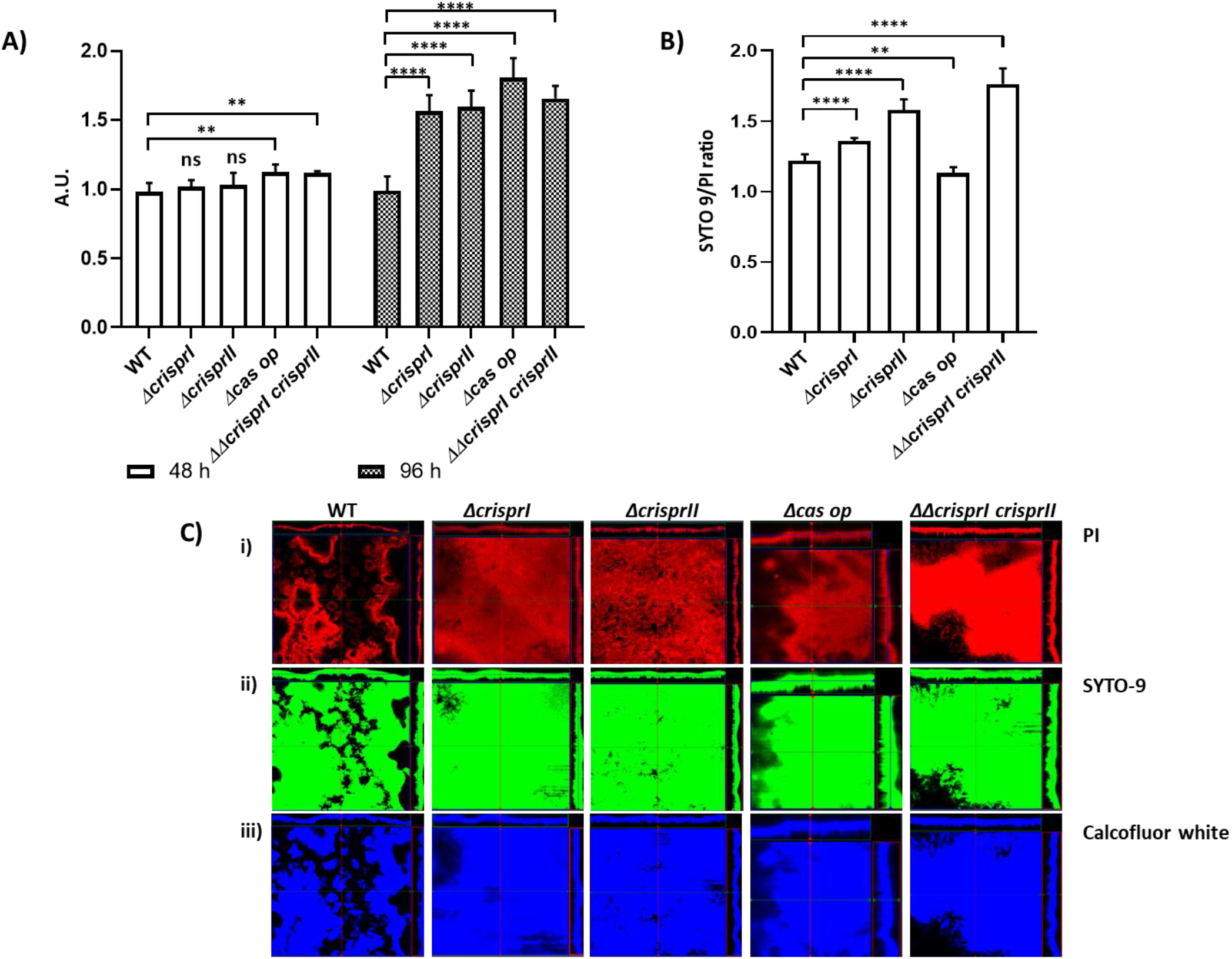
The CRISPR-Cas knockout strains had increased bacterial biomass (A), and SYTO 9/ PI ratio(B). **A.** The *S.* Typhimurium strain 14028s wildtype (WT), CRISPR (*ΔcrisprI, ΔcrisprII,* and *ΔΔcrisprI crisprII*) and *cas operon (Δcas op*) knockout strains were cultured in LB without NaCl media for different periods (48 h, and 96 h) at 25°C, static condition. The biomass of the strains was estimated by quantifying the dry weight of pellicle biofilms harvested post 48 h and 96 h incubations. The graph represents the dry pellicle weight (in gms) of each strain normalized by the dry pellicle weight (in gms) of WT at respective time points. **B-D.** The *S.* Typhimurium strain 14028s wildtype (WT), CRISPR (*ΔcrisprI, ΔcrisprII,* and *ΔΔcrisprI crisprII*) and *cas operon (Δcas op*) knockout strains were cultured in LB without NaCl media for 96 h, at 25°C, static condition. The pellicle biofilm formed was stained with SYTO 9, Propidium Iodide (PI), and Calcofluor white for 30 mins in the dark at RT. **B.** The graph represents ratio of mean intensity of SYTO 9 to mean intensity of PI, for respective strains. **C.** Orthogonal projections of CLSM images of wildtype and CRISPR-Cas knockout strains, stained with PI (i), SYTO 9 (ii), and Calcofluor white (iii). An unpaired t-test was used to determine significant differences between the WT and knockout strains. Error bars indicate SD. Statistical significance: *≤ 0.05, **≤ 0.01, ***≤ 0.001, ****<0.0001, ns = not significant. A.U., arbitrary units.

We next estimated the net content of the extracellular polymeric substances like proteins, DNA, and polysaccharides that comprise the ECM. The pellicle biofilms of all the knockout strains had significantly higher polysaccharide concentrations than WT (Supplementary figure, Fig. S6C). Similarly, the protein concentrations were significantly high in the pellicle biofilms of all the knockout strains except in *ΔΔcrisprI crisprII* (Supplementary figure, Fig. S6D). The DNA content was significantly higher only in the pellicle biofilm of *ΔcrisprI* and *Δcas op.* (Supplementary figure, Fig. S6E).

We further evaluated the expression of individual biofilm components like Curli and cellulose. Curli, thin aggregative fimbriae aid surface adhesion and provide cell-cell interaction while framing the biofilm architecture [19]. Less Curli production could also be one of the reasons for reduced ring biofilm formation by the knockout strains. Thus, we assessed the Curli production using whole-cell Congo red (CR) depletion assay for planktonic culture and pellicle biofilm. The CR depletion was less for both the planktonic culture (Supplementary figure, Fig. S8A) and pellicle biofilm (Fig.5A) of all the knockout strains, suggesting low levels of Curli protein. The results were further validated using an amyloid-specific indicator dye Thioflavin-T (ThT) [20]. The results confirm that the Curli production is less in all the four knockout strains (Fig. 5B). The cellulose production in the biofilm pellicle of all the strains was estimated by quantifying cellulose dry-weight at 48 h and 96 h. Interestingly, the cellulose content in the pellicles of the knockout strains was marginally lesser than that of WT at 48 h, but at 96 h the cellulose content was considerably higher (Fig.5C). The above-observed results for quantitative analysis of cellulose at 96 h were also substantiated by calcofluor binding assay (Fig.5D) and CLSM with calcofluor staining (Supplementary figure, Fig. S7C).

**Figure 5:**
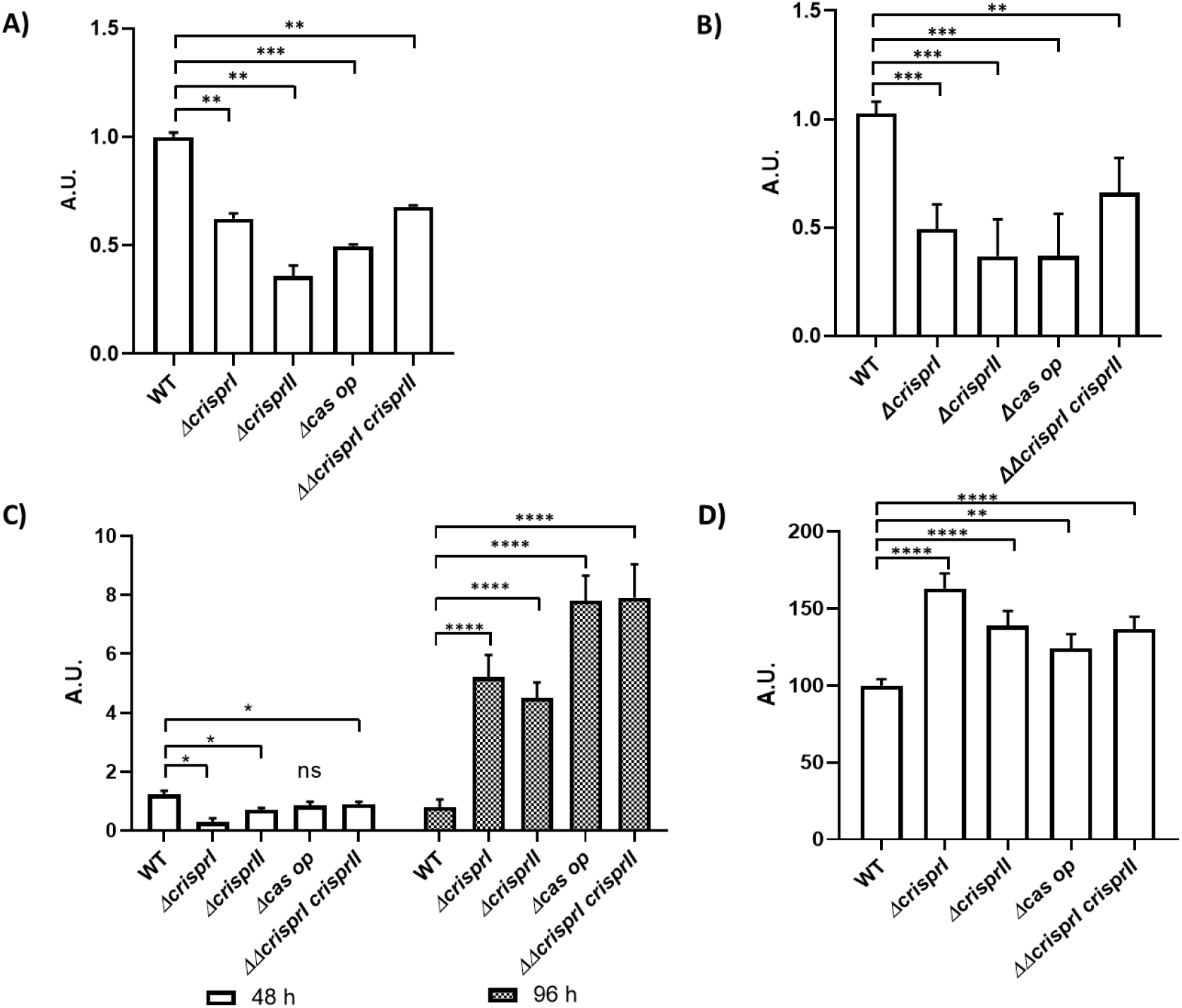
Pellicle biofilms of the CRISPR-Cas knockout strains showed variations in the productions of key components like curli (A & B), and cellulose (C & D). Curli production in the pellicle biofilms of wildtype, CRISPR, and *cas operon* knockout strains was assessed with the help of Congo red depletion (**A**), and Thioflavin (ThT) Fluorescence intensity (**B**). The *S.* Typhimurium strain 14028s wildtype (WT), CRISPR (*ΔcrisprI, ΔcrisprII,* and *ΔΔcrisprI crisprII*) and *cas operon (Δcas op*) knockout strains were cultured in LB without NaCl media 48 h, at 25°C, static condition. **A.** Congo red depletion was determined by measuring absorbance of bound Congo red at 500 nm. The graph represents normalized absorbance with respect to WT. **B.** Thioflavin (ThT) Fluorescence intensity was determined by measuring absorbance at excitation 440 nm and emission 482 nm. *Δcsg*D was used as a negative control. The graph represents intensity readings of each strain, normalized by intensity readings of WT. **C.** Cellulose production in the pellicle biofilms of wildtype, CRISPR, and *cas operon* knockout strains was quantitatively assessed by determining the cellulose dry weight in the pellicle biofilm. **D.** Qualitative analysis of amount of cellulose present in the pellicle-biofilm (96 h) was done by measuring the calcofluor bound, at excitation of 350 nm and emission 475 nm. The *S.* Typhimurium strain 14028s wildtype (WT), CRISPR (*ΔcrisprI, ΔcrisprII* and *ΔΔcrisprI crisprII*) and *cas operon (Δcas op.*) knockout strains were cultured in LB without NaCl media 96 h, at 25°C, static condition. An unpaired t-test was used to determine significant differences between the WT and knockout strains. Error bar indicates SD. Statistical significance: *≤ 0.05, **≤ 0.01, ***≤ 0.001, ****<0.0001, ns = not significant. A.U., arbitrary units.

Curli content in the pellicle biofilm is related to surface elasticity, thereby providing mechanical strength to the biofilm [21]. As Curli protein was lesser in pellicles of knockout strains, we determined the pellicle biofilm strength using a glass bead assay [21]. The pellicles of the knockout strains were easily disrupted with lesser weight while enduring ~50% less weight than the WT pellicles could sustain (Supplementary figure, Fig. S8C). The results suggest that knockout strains' pellicles are weaker due to lesser Curli production.

### The CRISPR-Cas knockout strains show altered expression of biofilm-related genes

To understand the temporal variations in biofilm formation by the CRISPR-Cas knockout strains, we checked the regulation of biofilm-related genes using RT-PCR. We first assessed the expression of genes governing motility, like *fliC* (flagellin subunit), *flgK* (hook protein), *yddX* (biofilm modulation protein, controlling regulatory pathway of flagellar assembly), and *flgJ* (peptidoglycan hydrolyzing flagellar protein). All the knockout strains showed reduced expression of these genes (Fig.6A), except *flgJ* (Supplementary figure, Fig. S9A). Next, to comprehend the observed variations in the LPS profile of the knockout strains (Supplementary figure, Fig. S5), we analyzed the expression of a few representative LPS genes within *rfa* (LPS core synthesis) and *rfb* (O-antigen synthesis) gene clusters. The *rfaC* (lipopolysaccharide heptosyltransferase I), and *rfbG* (DP-glucose 4,6-dehydratase) genes were upregulated in all the knockout strains (Fig.6B), whereas *rfbI,* coding for core LPS region was downregulated in all the knockout strains except in *ΔΔcas op.* (Supplementary figure, Fig. S9B).

**Figure 6:**
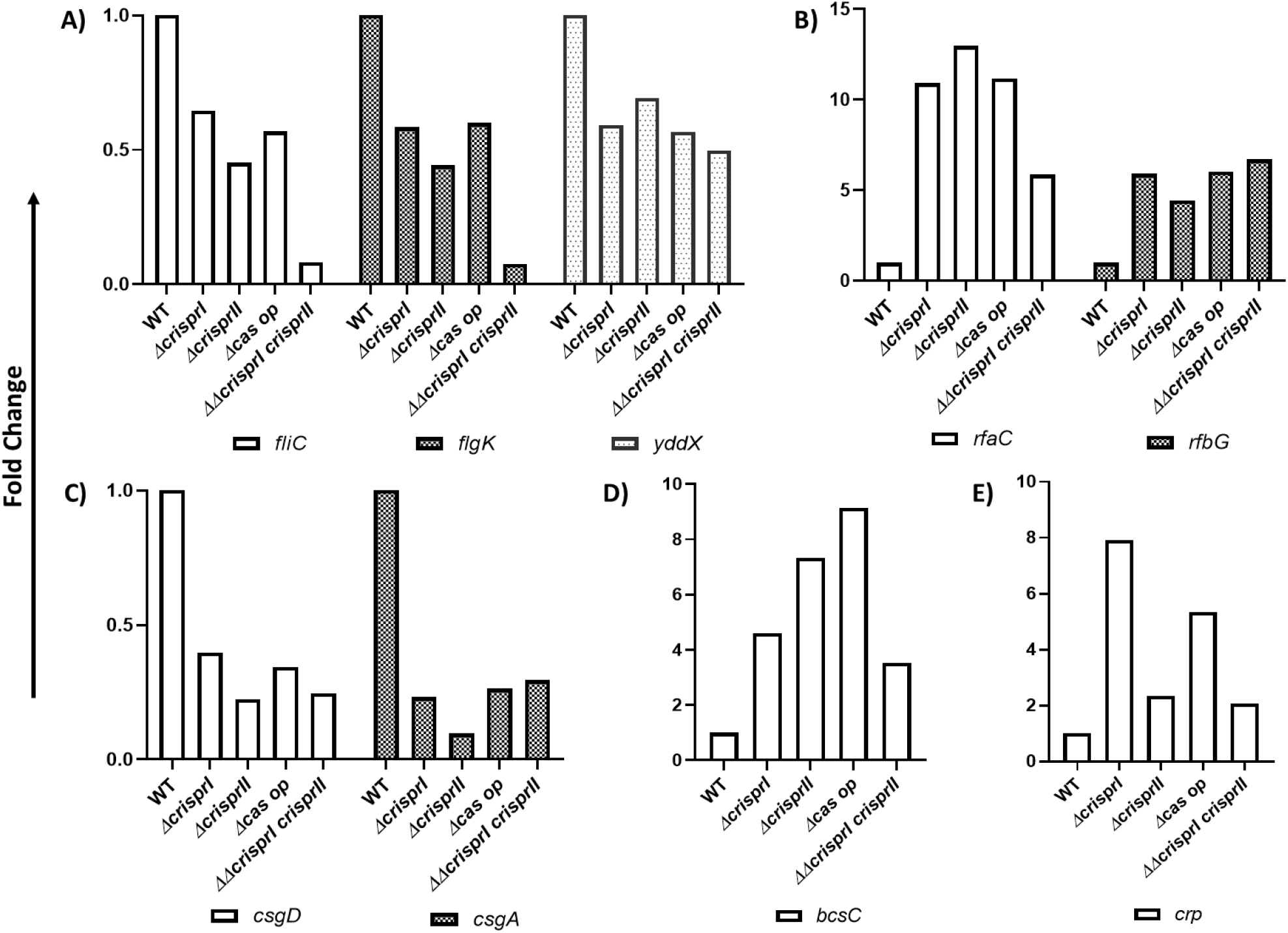
CRISPR-Cas system knockout strains showed differences in the expressions of genes associated with flagella (A), the production of LPS (B), Curli (C), cellulose (D), and cAMP regulated protein (*crp*) (E) when compared to WT. The *S.* Typhimurium strain 14028s wildtype (WT), CRISPR (*ΔcrisprI, ΔcrisprII* and *ΔcrisprI ΔcrisprII*) and *cas operon (Δcas op*) knockout strains were cultured in LB without NaCl media for 24 h, at 25°C, static condition. Total RNA was isolated from bacteria using TRIzol reagent as per the manufacturer's instructions. 1 μg of RNA was used for cDNA synthesis, followed by qRT-PCR. Relative expression of the gene was calculated using the 2^-ΔΔCt^ method, and normalized to reference gene *rpoD.*

The *csgA* gene responsible for producing the Curli fibers was downregulated in knockout strains (Fig.6C). The expression of *csgA* is controlled by the master regulator *csgD*, which too had reduced expression in the knockout strains (Fig.6C). The expression of *crp* gene encoding for cAMP regulating protein, a *csgD* repressor [22], was high in the knockout strains (Fig.6E). *CsgD* also controls the expression of cellulose synthesis genes (*bcsABZC*). Notably, the expression of *bcsA* (cellulose synthase catalytic subunit A) was only marginally low in the knockout strains (Supplementary figure, Fig. S9C) but *bcsC* (subunit involved in the export of cellulose to extracellular matrix [23]) was 2-fold upregulated (Fig.6D). The observed results hint at *csgD* independent regulation [24] of *bcsC* in the knockout strains.

## Discussion

Biofilm formation in *Salmonella* is finely regulated, helping the bacteria to sustain various environmental insults while aiding in its persistence within and outside the host [25]. Recently, the CRISPR-Cas system has been implicated to play a role in endogenous gene regulation [5] and biofilm formation in various bacteria, including *Salmonella* [6], [7]. Cui *et al.* demonstrated that Cas3 positively regulates biofilm formation in *S. enterica* subsp*. enterica* ser. Enteritidis [6]. However, our study determined that the Cas proteins negatively regulate biofilm formation in *S.* Typhimurium. This discrepancy in the results could be related to the differences in CRISPR spacers within these serovars [26] or differences in *cas* gene expression observed in both studies. The *cas* genes were upregulated in the *cas3* mutant strain of serovar Enteritidis. This implies that the increased expression of Cas proteins (except Cas3) could have suppressed biofilm in serovar Enteritidis. While in our study, the entire operon was deleted, thereby no *cas* gene expression and hence enhanced biofilm formation. Furthermore, our study also demonstrated that CRISPR-I and CRISPR-II arrays negatively regulated pellicle biofilm formation in *S.*Typhimurium. Correspondingly, a study by Medina *et al.* suggests that the CRISPR-Cas system suppresses the surface-biofilm formation (at 24 h) in *S*. Typhi [7]. Intriguingly, we found that the CRISPR-Cas system of *S*. Typhimurium positively regulates surface-biofilm while repressing pellicle-biofilm. We speculate that the difference in our data on surface-biofilm and that of Medina *et al.* could be because serovar Typhi and Typhimurium differ in arrangement and sequence of *cas* genes, as well as in the CRISPR-I array [6], [7]. Could the differential evolution of the CRISPR-Cas system possibly be the reason for the two serovars' distinct biofilm phenotypes? Or could it be due to differences in the CRISPR spacers? These deductions need further exploration.

We next explored the underlying mechanisms of biofilm regulation by the CRISPR-Cas system. Biofilm formation is a complex mechanism requiring coordination between multiple factors and processes. Flagellar motility is essential for cell-cell adhesion and forming microcolonies at the initial stages [16]. Our study showed that the CRISPR-Cas knockout strains are less motile, thereby explaining lesser biofilm formation at 24 h by CRISPR-Cas knockout strains. Nevertheless, as the biofilm progresses, the requirement of flagella becomes negligible, and its expression is repressed [27]. In accordance, we found that FliC expression was absent in pellicle biofilms of all the strains at 96 h. The FliC subunit is also crucial for cholesterol binding and biofilm development on gallstones [28], [29]. The decreased biofilm formation by the CRISPR-Cas knockout strains in tube biofilm assay could be attributed to decreased FliC expression. The reduction in FliC expression is also reflected in reduced swarming motility of the knockout strains, but it is not proportionate to the observed trend in FliC expression. This disparity could be due to variation in the LPS that acts as a wettability factor favoring swarming while inhibiting biofilm formation [30]. Interestingly, our study displayed such a relation; all the knockout strains showed reduced swarming but enhanced biofilm formation. Further, despite showing minimal FliC expression amongst all knockout strains, *ΔΔcrisprI crisprII* had considerable swarming motility.

The CRISPR-Cas knockout strains had altered LPS profile with a difference in the LPS gene expression. The *rfaC* (part of *rfa* gene cluster: responsible for LPS core synthesis), and *rfbG* (part of *rfb* gene cluster: responsible for O-antigen synthesis) genes were upregulated in the knockout strains. At the same time, *rfbI* was significantly downregulated only in *ΔΔcrisprI crisprII*. Besides, studies suggest the plausible conversion of LPS to exopolysaccharides that contribute to external slime [31]. The increased exopolysaccharides in the pellicle of CRISPR-Cas knockout strains may also be attributed to this, along with the observed increase in cellulose production. The pellicles formed by the CRISPR-Cas knockout strains are thicker (owing to more bacterial mass and EPS secretion[16]) than that of the WT, confirming the formation of multilayered pellicle biofilms, as evidenced by SEM and CLSM analysis. As per SEM analysis, the air-exposed pellicle biofilm architecture of *ΔcrisprII* and *ΔΔcrisprI crisprII* appears similar, indicating that *crisprII* could act upstream of *crisprI*. This observation is seconded by our LPS profiling data, where the banding pattern of *ΔcrisprII* and *ΔΔcrisprI crisprII* are similar.

The EPS overproducing variants reportedly have rough and wrinkled biofilm [32]. This supports our observation that the CRISPR-Cas knockout strains overproduce EPS and display intricate wrinkled patterns in the pellicle biofilm. These wrinkled patterns appeared fractal-like (Supplementary figure, Fig. S9B), as reported in *Vibrio cholerae* [33]. Such morphology could aid bacterial growth of the CRISPR-Cas knockout strains due to the increased surface area that presumably facilitates the nutrient supply [33]. Consistently, the bacterial mass was higher in the knockout strains with more viable bacteria, as evidenced by the resazurin assay and SYTO9-PI staining.

The ECM scaffold of pellicle biofilm majorly comprises cellulose and Curli that define the long-range and short-range interactions, respectively, thereby providing mechanical integrity[34]. The pellicle biofilms of CRISPR-Cas knockout strains have higher cellulose but lesser curli content. This could probably be the reason for the weaker pellicle biofilm of the CRISPR-Cas knockout strains that quickly collapsed in the glass bead assay. Further, high cellulose in the pellicles of the CRISPR-Cas knockout strains means high water retention that can hamper intermolecular forces in the matrix by decreasing the hydrogen bond interactions. Additionally, less Curli could lead to low tensile strength of the pellicle biofilm of the CRISPR-Cas knockout strains. Higher cellulose and lesser Curli could also explain reduced surface-biofilm (ring biofilm at 24 and 96 h) in the CRISPR-Cas knockout strains. High cellulose may inhibit the formation of surface-biofilm as it can coat the curli fibers required for surface attachment[35]. Though the cellulose content was high in pellicles of the CRISPR-Cas knockout strains, the expression of cellulose synthase, *bcsA*, was unaltered; indeed it was marginally low in all the knockout strains. Besides, the intracellular cellulose concentration in the CRISPR-Cas knockout strains was less than the WT (as estimated using anthrone assay, supplementary figure, Fig. S8B). This could be explained through the upregulated *bcsC*, encoding an exporter of cellulose subunits that could export cellulose units to the extracellular milieu[23]. We hypothesized that this secreted cellulose is quickly incorporated in the pellicle, increasing cellulose content in the pellicles of the knockout strains.

Apart from reduced expression of *csgA* and marginal repression of *bcsA*, we found that *csgD*, the activator of *csgBAC* and *bcsABZC* was also downregulated in the knockout strains. In order to gain mechanistic insight into the CRISPR-Cas mediated biofilm regulation, we checked the expression of the further upstream regulator, CRP. CRP negatively regulates *csgD* in *S*. Typhimurium[36]. The expression of *crp* was significantly upregulated in the knockout strains, which explains the repression of *csg* and *bcsA*. The conflicting upregulation of *bcsC*, the last gene of *bcsABZC*, could be through the crRNA binding to the *bcsC* gene. The CRISPR spacers (spacer 11, 15, and 19 of CRISPR1 array and spacer 18 and 26 of CRISPR2 array) show partial complementarity to the *bcsC* gene (Supplementary figure, Fig. S11) and hence could regulate the expression of *bcsC*. CRP also activates *flhDC,* a flagellar master operon[37] that further activates the expression of class 2 genes, including *fliA.* The *fliA* gene encodes the flagellar-specific transcription factor σ^28^, which directs the expression of class 3 genes like *fliC* and *flgK*. Before the assembly of hook-basal body structure, it is held inactive by the anti-σ^28^ factor, *flgM*[38]. YddX, a BDM homolog (Biofilm-dependent modulation protein) interacts with FlgM to repress its function as an anti-σ^28^ factor[39]. Our study observed a significant downregulation of *yddX* in the knockout strains. Low YddX would mean that FlgM would sequester σ^28^, inhibiting the transcription and expression of class 3 genes, including *fliC* and *flgK.* This explains the impaired motility of the CRISPR-Cas knockout strains.

In a nutshell, CRISPR-Cas system facilitates surface-attached biofilm formation while repressing the pellicle biofilms by acting on different biofilm regulators. The mechanism is summarized in Fig. 7.

**Figure 7.**
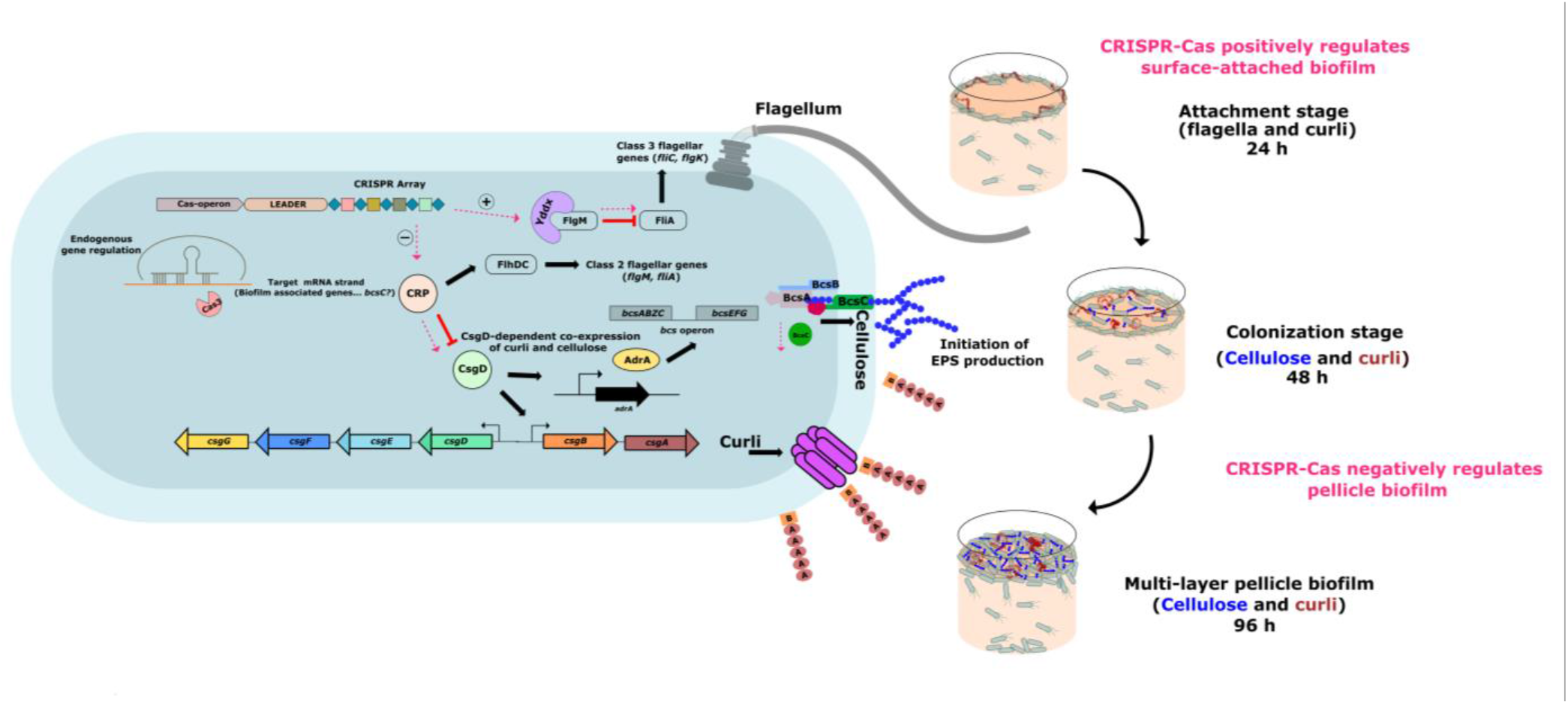
Differential regulation of surface-attached and pellicle-biofilm formation in *Salmonella* Typhimurium by the CRISPR-Cas system. The CRISPR-Cas system differentially regulates surface-attached and pellicle-biofilm formation via modulation (pink dotted lines) of biofilm-associated genes (*crp, yddx, and bcsC*). CRP acts on FlhDC, which further governs the expression of class 2 flagellar genes (*flgM* and *fliA*). FlgM inhibits FliA mediated expression of class 3 flagellar genes. Yddx relieves the inhibition of FliA by binding to FlgM, thereby inactivating it. We propose that CRISPR-Cas positively regulates *yddx*, whereby it sequesters FlgM and upregulates the expression of the flagellar subunit. CRP also inhibits CsgD, which in turn governs the production of Curli and cellulose. Our study suggests that the CRISPR-Cas system mediates the expression of CsgD by suppressing *crp* expression, and independently represses the expression of cellulose exporter, BcsC. Taken together, the CRISPR-Cas system enhances flagella and Curli production and hence surface-attached biofilm formation. Additionally, it suppresses cellulose export to the extracellular milieu, thus negatively regulating pellicle biofilm formation.

## Acknowledgments

This work was supported by the Department of Science and Technology, Science and Engineering Research Board (Grant No. ECR_2017_002053) to SAM.

## Supplementary information

### Material and Methods

#### Analyzing Lipopolysaccharide (LPS) profiles

LPS lysis buffer (2 mL of 20% SDS, 800 μL ß-Mercaptoethanol, 200 μL bromophenol, 2 mL glycerol, 15 mL of 1M Tris-HCl) was added to the pellicle biofilms and were rinsed twice with distilled water. The samples were then lysed using TissueLyser LT (QIAGEN, Germany) at 50 Hz for 10 mins. The lysates were heated at 100°C, 10 mins followed by DNase (1 μg/μL), RNase (20 μg/μL), and Proteinase-K (20 μg/μL) treatment. Crude LPS thus obtained was resolved using SDS–polyacrylamide gel electrophoresis (SDS-PAGE) with 15% separating gel. The LPS profile was detected using ProteoSilver Silver stain Kit (SIGMA-ALDRICH, USA).

#### Determining Pellicle Strength

The strains were cultured in LB without NaCl media for 96 h, at 25°C, static condition. The pellicle biofilm strength was determined by addition of glass beads (1 mm, HiMedia) using a tweezer until disruption (collapse of pellicle to the bottom). The weight of glass beads that collapsed the pellicle was recorded.

#### Quantification of extracellular matrix (ECM) components

The 96 h pellicle biofilms were washed with sterile water and sonicated on ice, 15 kHz for 30 secs. The samples were centrifuged and supernant was used for the analysis. The DNA and protein concentrations in the supernatants of each sample were estimated spectrophotometrically using BioSpectrometer^®^ basic (Eppendorf, Germany). The exopolysaccharides were quantified by the phenol-sulphuric acid method[1] followed by absorbance at 490 nm.

---

**Supplementary Table 1:** Bacterial strains used in this study

| Bacterial Strain | Genotype and Characteristics | Source/ref |
| --- | --- | --- |
| *Salmonella enterica* serovars Typhimurium 14028s | WT 14028s | A kind gift from Prof. Dipshikha Chakravorty, Indian Institute of Science, India |
| ΔcrisprI | WT 14028s ΔcrisprI::Chl (Chl<sup>r</sup>) | This study |
| ΔcrisprII | WT 14028s ΔcrisprII::Chl (Chl<sup>r</sup>) | This study |
| Δcas op | WT 14028s Δcas operon::Chl (Chl<sup>r</sup>) | This study |
| $\Delta\Delta$crisprI crisprII | WT 14028s ΔcrisprI::Kan :ΔcrisprI::Chl (Kan<sup>r</sup>, Chl<sup>r</sup>) | This study |
| ΔfliC | WT 14028s ΔfliC::Kan (Kan<sup>r</sup>) | Marathe *et al.*, 2016 |
| ΔcsgD | WT 14028s *csgD* : : Chl (Chl<sup>r</sup>) | A kind gift from Prof. Dipshikha Chakravorty, Indian Institute of Science, India |
| WT60 | WT 14028s transformed with empty pQE60 vector | This study |
| ΔcrisprI 60 | ΔcrisprI transformed with empty pQE60 vector | This study |
| ΔcrisprII 60 | ΔcrisprII transformed with empty pQE60 vector | This study |
| Δcas op 60 | Δcas op transformed with empty pQE60 vector | This study |
| $\Delta\Delta$crisprI crisprII 60 | $\Delta\Delta$crisprI crisprII transformed with empty pQE60 vector | This study |
| ΔcrisprI +pcrisprI | ΔcrisprI complemented with functional CRISPR I array cloned in pQE60 | This study |
| ΔcrisprII +pcrisprII | ΔcrisprII complemented with functional CRISPR II array cloned in pQE60 | This study |

**Supplementary Table 2:**
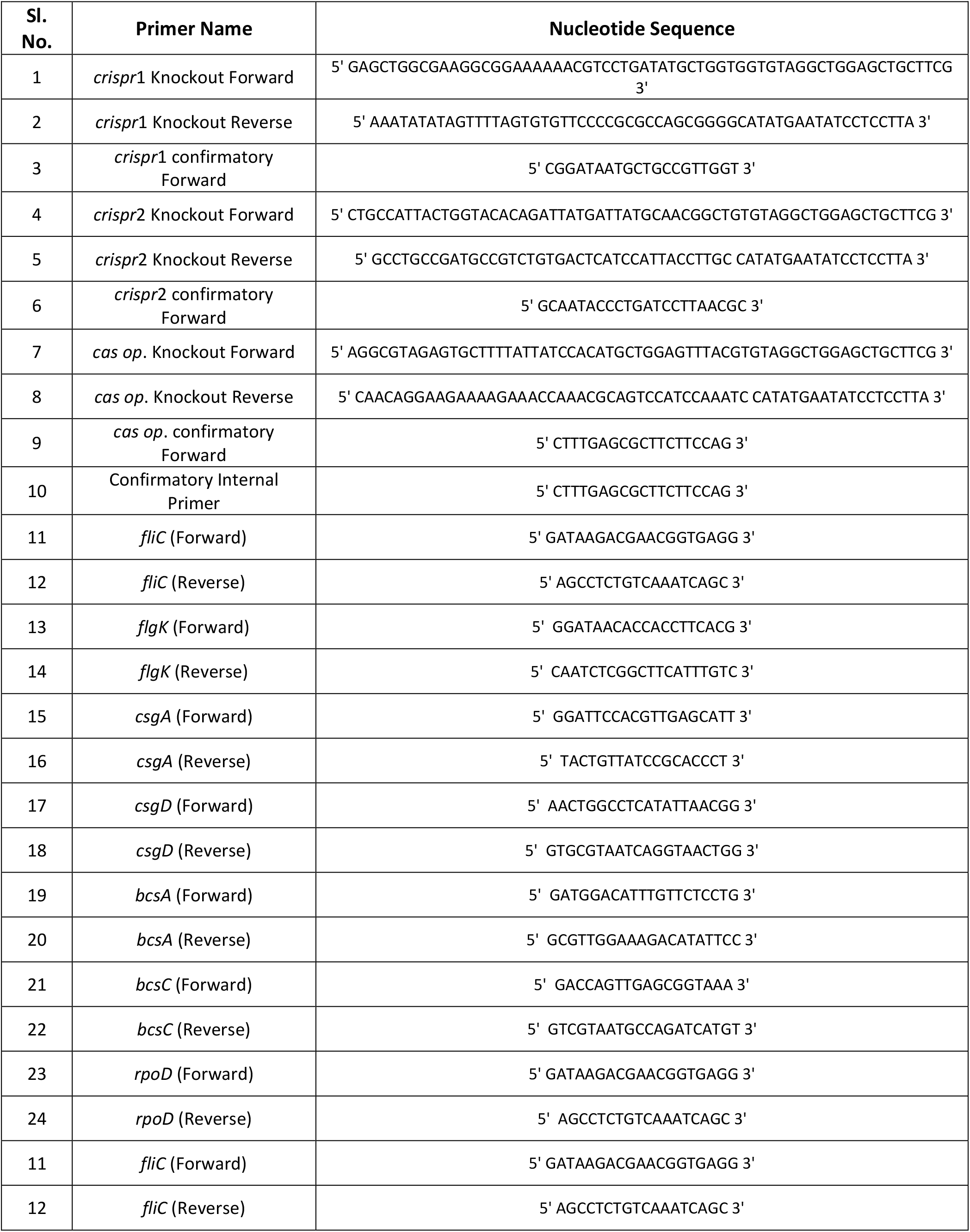
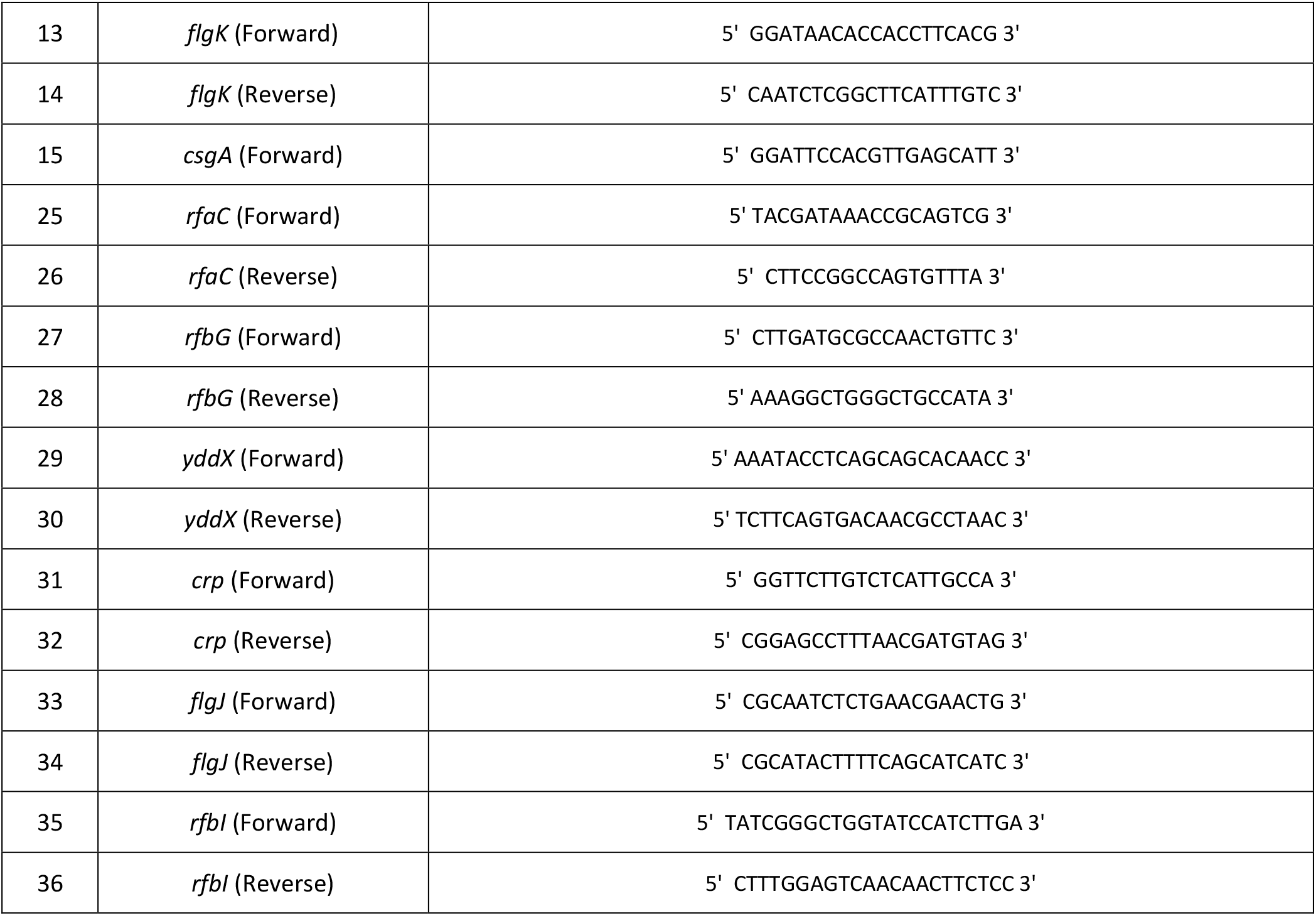
Primers used in this study

**Supplementary Figure S1:**
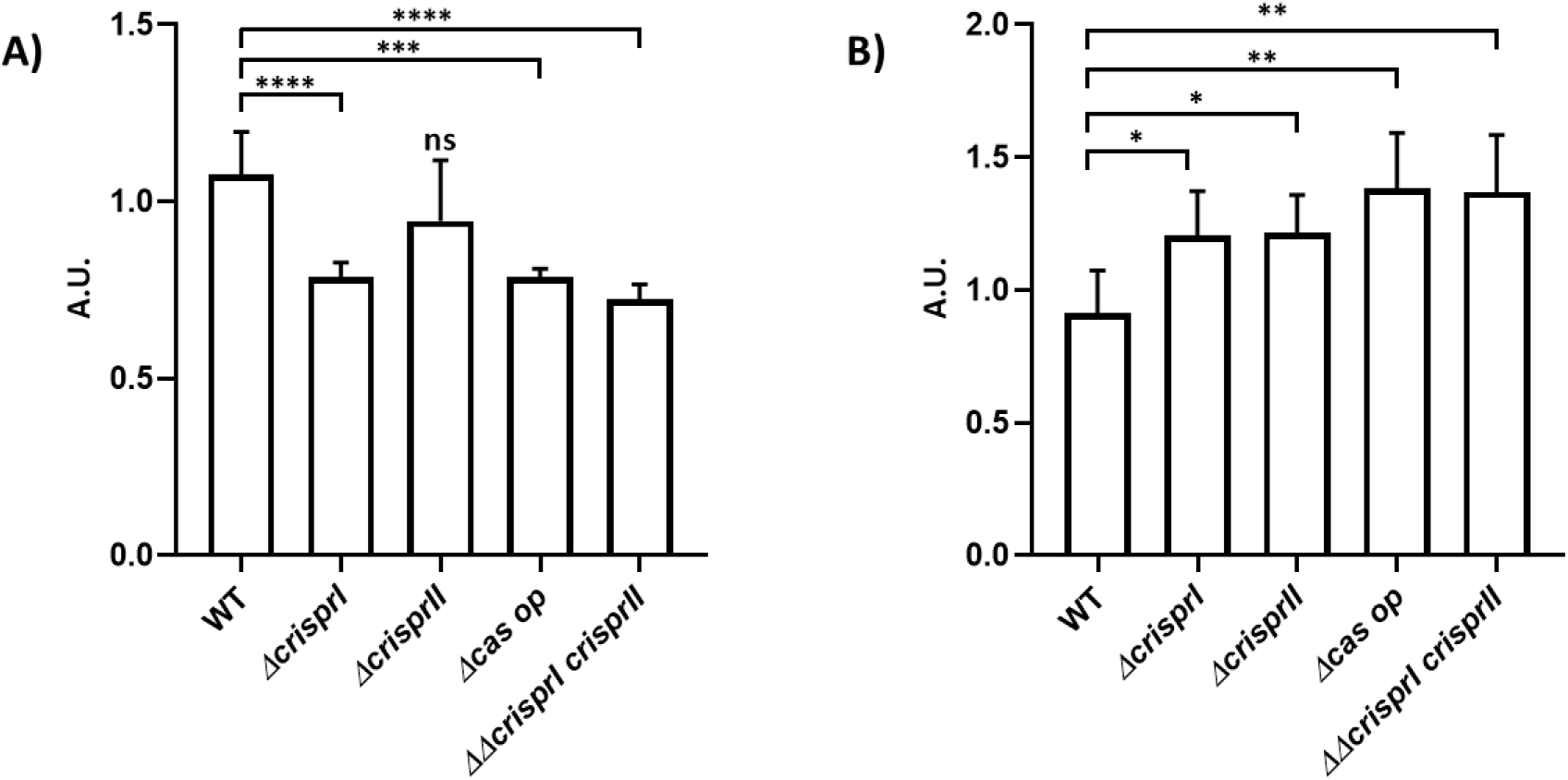
The CRISPR-Cas system knockout strains of *S. enterica* subsp. *enterica* serovar Typhimurium 14028s showed reduced biofilm formation at the solid-liquid interface (A), while these strains showed increased floating biofilm (pellicle) (B). The *S.* Typhimurium strain 14028s wildtype (WT), CRISPR (*ΔcrisprI, ΔcrisprII,* and *ΔΔcrisprI crisprII*) and *cas operon (Δcas op*) knockout strains were cultured in Tryptic Soy Broth (TSB) media for 96 h, at 25°C, static condition in 24-well plastic plate. The biofilm formation was estimated using the crystal violet staining method. The graph represents OD_570nm_ for each strain, normalized by OD_570nm_ of WT biofilm. An unpaired t-test was used to determine significant differences between the WT and knockout strains. Error bars indicate SD. Statistical significance: *≤ 0.05, **≤ 0.01, ***≤ 0.001, ****<0.0001, ns = not significant. A.U., arbitrary units.

**Supplementary Figure S2:**
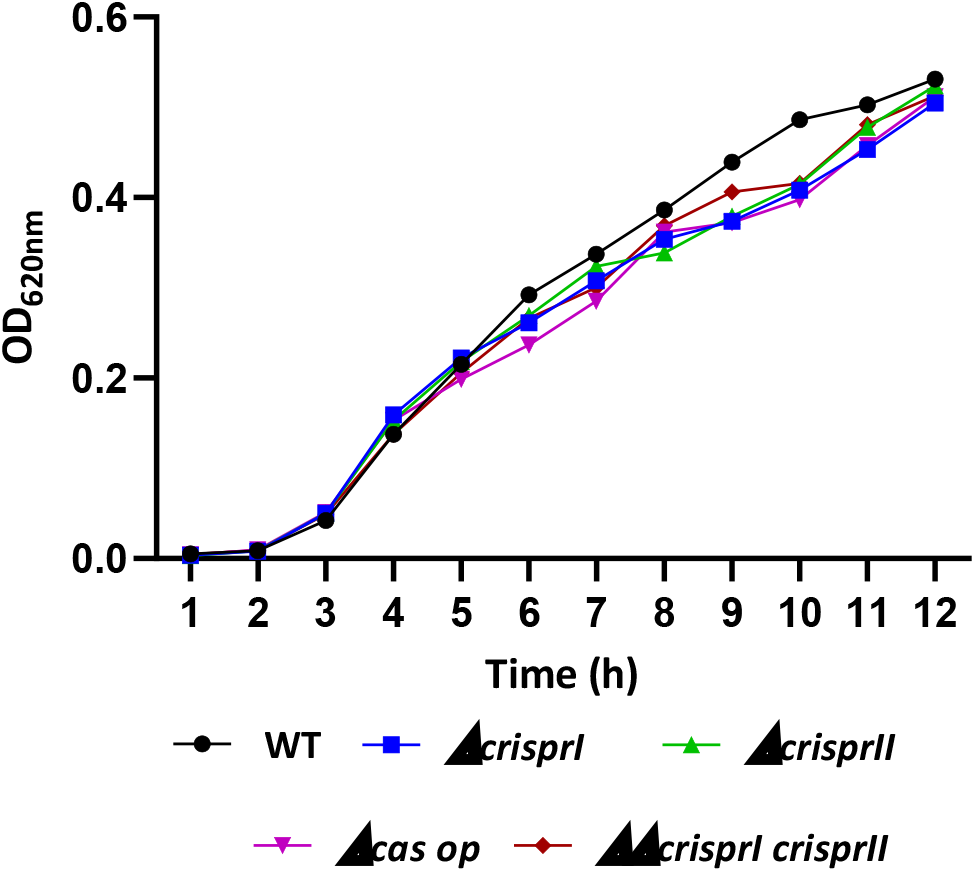
The CRISPR-Cas system knockout strains of *S. enterica* subsp. *enterica* serovar Typhimurium 14028s showed a similar growth trend to wildtype in LB without NaCl media. The *S.* Typhimurium strain 14028s wildtype (WT), CRISPR (*ΔcrisprI, ΔcrisprII,* and *ΔΔcrisprI crisprII*) and *cas operon (Δcas op*) knockout strains were cultured in LB without NaCl media for 12 h, at 37°C, shaking condition. The graph represents OD_620nm_ against time (h) for each strain.

**Supplementary Figure S3:**
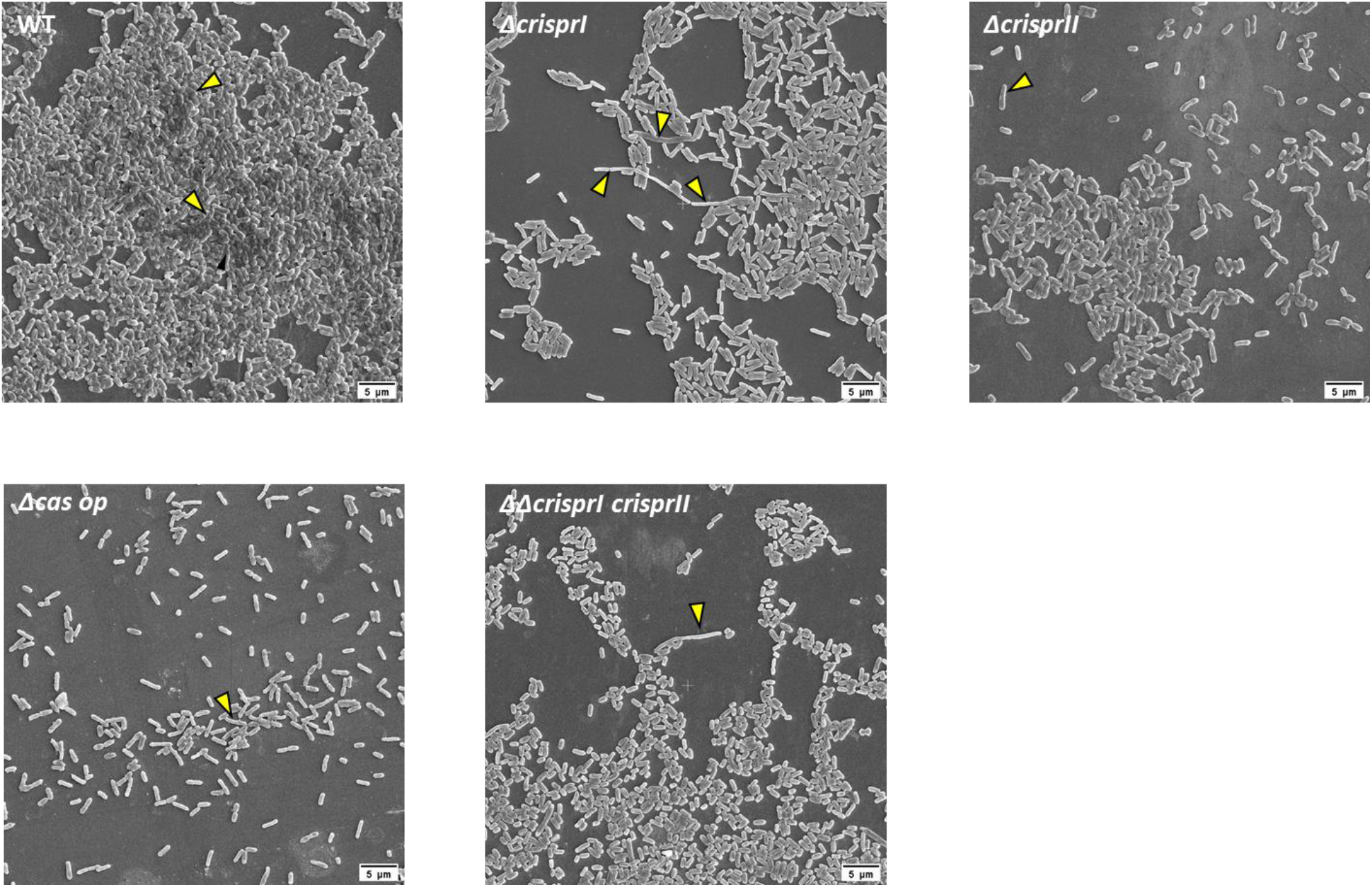
Morphology of air-exposed side of pellicle biofilm at early (24 h) time point. The knockout (*ΔcrisprI, ΔcrisprII, Δcas op.,* and *ΔΔcrisprI crisprII*) strains formed patchy bacterial aggregates, in comparison to wildtype (WT). WT biofilm had tightly packed bacterial aggregates covering a larger area, with a few dome-like structures (arrow-head in the WT micrograph). Few elongated cells (arrow-head in the micrographs) were also observed in the biofilms of the knockout strains. The strains were grown in LB without NaCl media for 24 h, at 25°C, static conditions. The pellicle biofilms formed were fixed using 2.5% glutaraldehyde and dehydrated with increasing ethanol concentrations. The images were captured at 5000X magnification and scaled to bar.

**Supplementary Figure S4:**
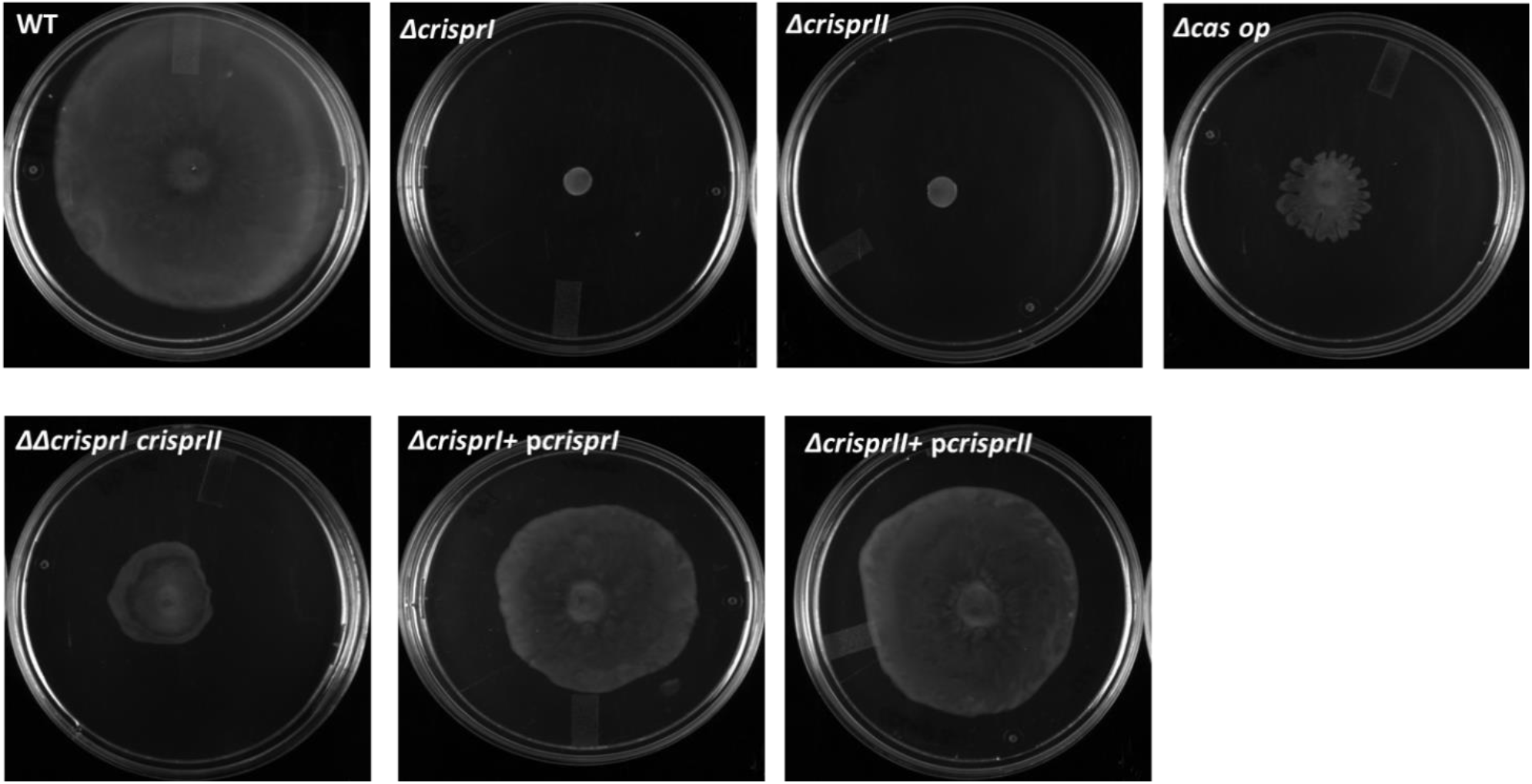
CRISPR-Cas system knockout strains show reduced swarming motility. Swarm plates (0.5% agar, 20g/L of LB and 0.5% glucose) were point inoculated with overnight cultures and incubated at 37°C for 9 h. The complement strains (*ΔcrisprI* + p*crisprI, ΔcrisprII*+ p*crisprII*) showed reversal of swarming ability confirming the mutation process was not polar.

**Supplementary Figure S5:**
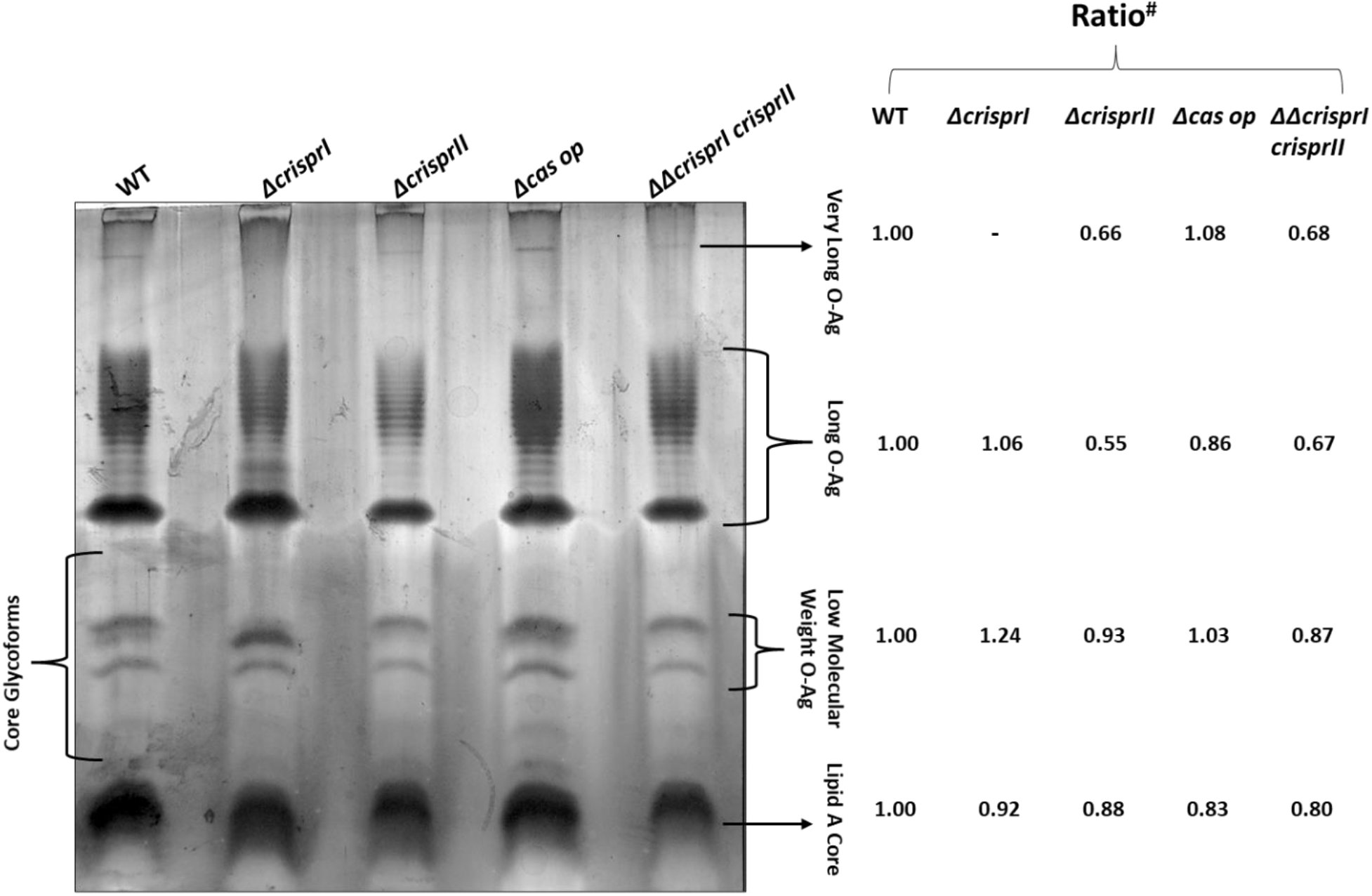
Silver-stained Lipopolysaccharide (LPS) profiling of wildtype (WT), and CRISPR-Cas system knockout strains. The variation in O-antigen was analyzed by LPS profiling. The strains were grown in LB without NaCl media for 96 h, at 25°C, static conditions. Pellicle biofilm was homogenized and heated, followed by DNase, RNases, and Proteinase-K treatment to extract crude LPS. The processed samples were loaded on 15% SDS-PAGE MIDI gel, which was later stained using a silver staining kit. Variations in banding pattern and intensity between knockout (*ΔcrisprI, ΔcrisprII, Δcas op,* and *ΔΔcrisprI crisprII*) strains and WT were observed in long O-Ag, low molecular weight O-Ag, and core glycoforms regions. #Ratio indicates the intensity of the bands observed on the gel for all strains normalized by the corresponding band's intensity of wildtype sample.

**Supplementary Figure S6:**
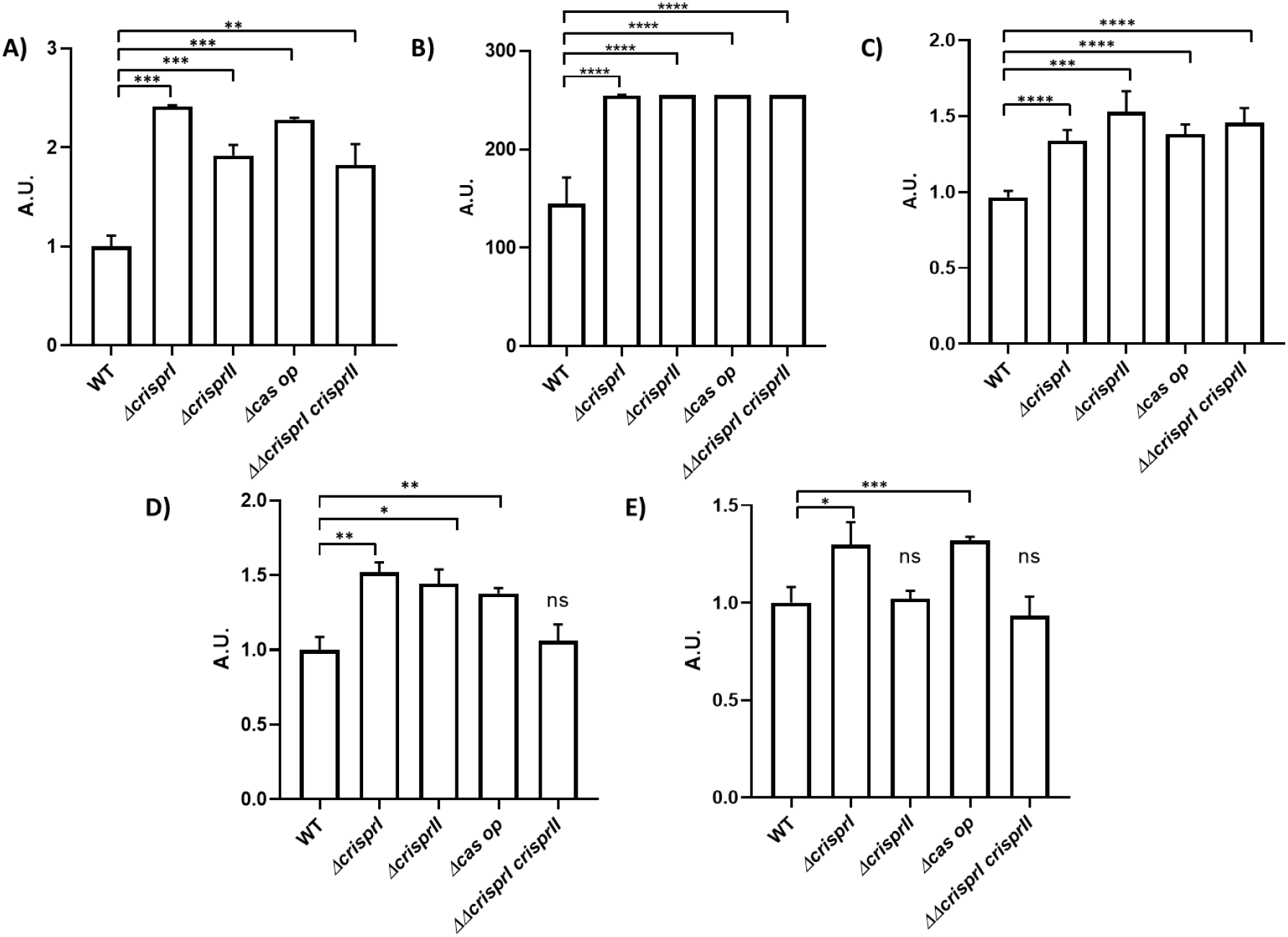
Compared to WT, CRISPR-Cas system knockout strains show differences in their metabolic activity (A), bacterial cell population (B), and ECM components like polysaccharides (C), protein (D), and DNA (E). **A.** The metabolic activity was assessed by resazurin assay. *S.* Typhimurium strain 14028s wildtype (WT), CRISPR (*ΔcrisprI, ΔcrisprII* and *ΔΔcrisprI crisprII*) and *cas operon (Δcas op*) knockout strains were cultured in LB without NaCl media for 96 h, at 25°C, static condition. The pellicle biofilm formed after 96 h incubation was stained with resazurin dye and fluorescence was measured using a fluorimeter at excitation (λ_Ex_) 550 nm and emission (λ_Em_) of 600 nm. The graph represents the fluorescence intensity observed for each strain normalized by the fluorescence intensity of WT. **B** The pellicle biofilms formed by all the strains were stained with SYTO 9, for 30 mins in the dark, at RT. The graph represents the mean intensity of SYTO9 observed for each strain. **C.** The exopolysaccharides from the pellicle biofilms were quantified by the phenol-sulfuric acid method, by measuring absorbance at 490 nm. The graph represents absorbance observed at 490 nm for each strain normalized by absorbance observed at 490 nm for the WT sample. **D & E.** The protein and DNA concentrations in each sample were estimated spectrophotometrically and were further normalized by absorbance for WT in each case. An unpaired t-test was used to determine significant differences between the WT and knockout strains. Error bars indicate SD. Statistical significance: *≤ 0.05, **≤ 0.01, ***≤ 0.001, ****<0.0001, ns = not significant. A.U., arbitrary units.

**Supplementary Figure S7.**
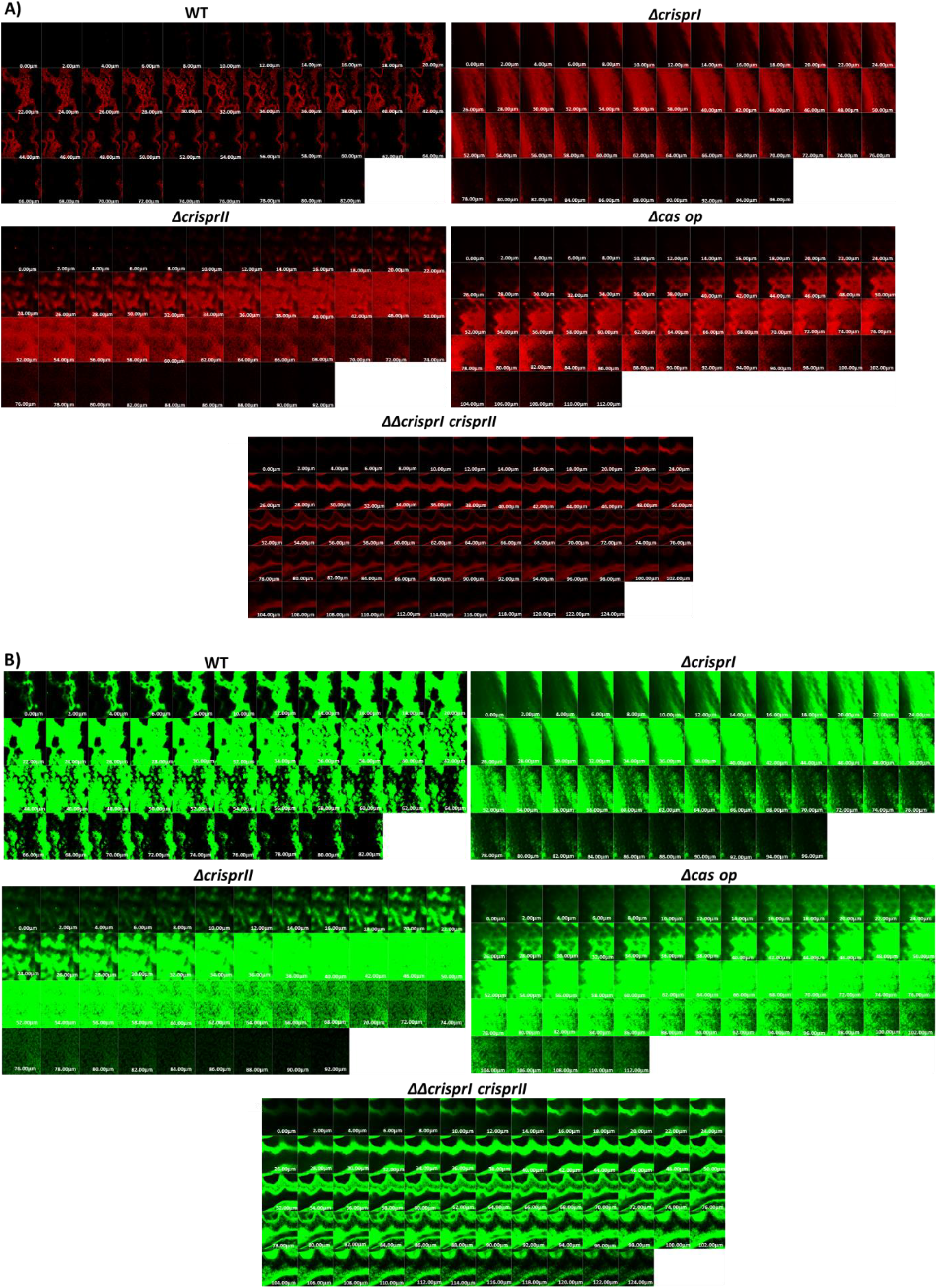
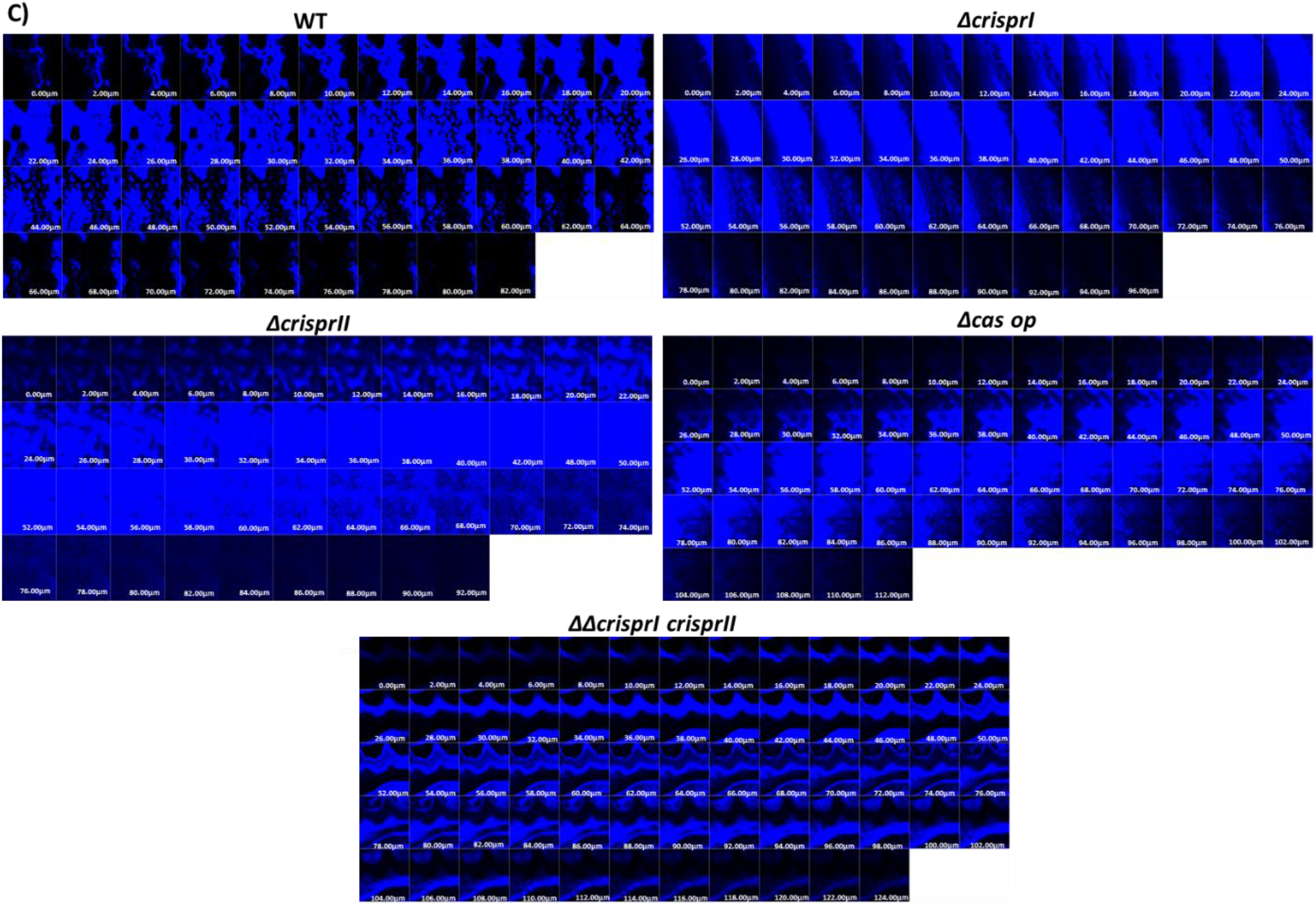
CLSM images (stacks) of wildtype and CRISPR-Cas knockout strains stained with Propidium Iodide (A), SYTO 9 (B) and Calcofluor white (C). The *S.* Typhimurium strain 14028s wildtype (WT), CRISPR (*ΔcrisprI, ΔcrisprII,* and *ΔΔcrisprI crisprII*) and *cas operon (Δcas op*) knockout strains were cultured in LB without NaCl media for 96 h, at 25°C, static condition. The pellicle biofilm formed was stained with Propidium Iodide (PI), SYTO 9, and Calcofluor white for 30 mins in the dark, at RT. The Z-stacks of the CLSM images were captured and the stacks are represented here.

**Supplementary Figure S8:**
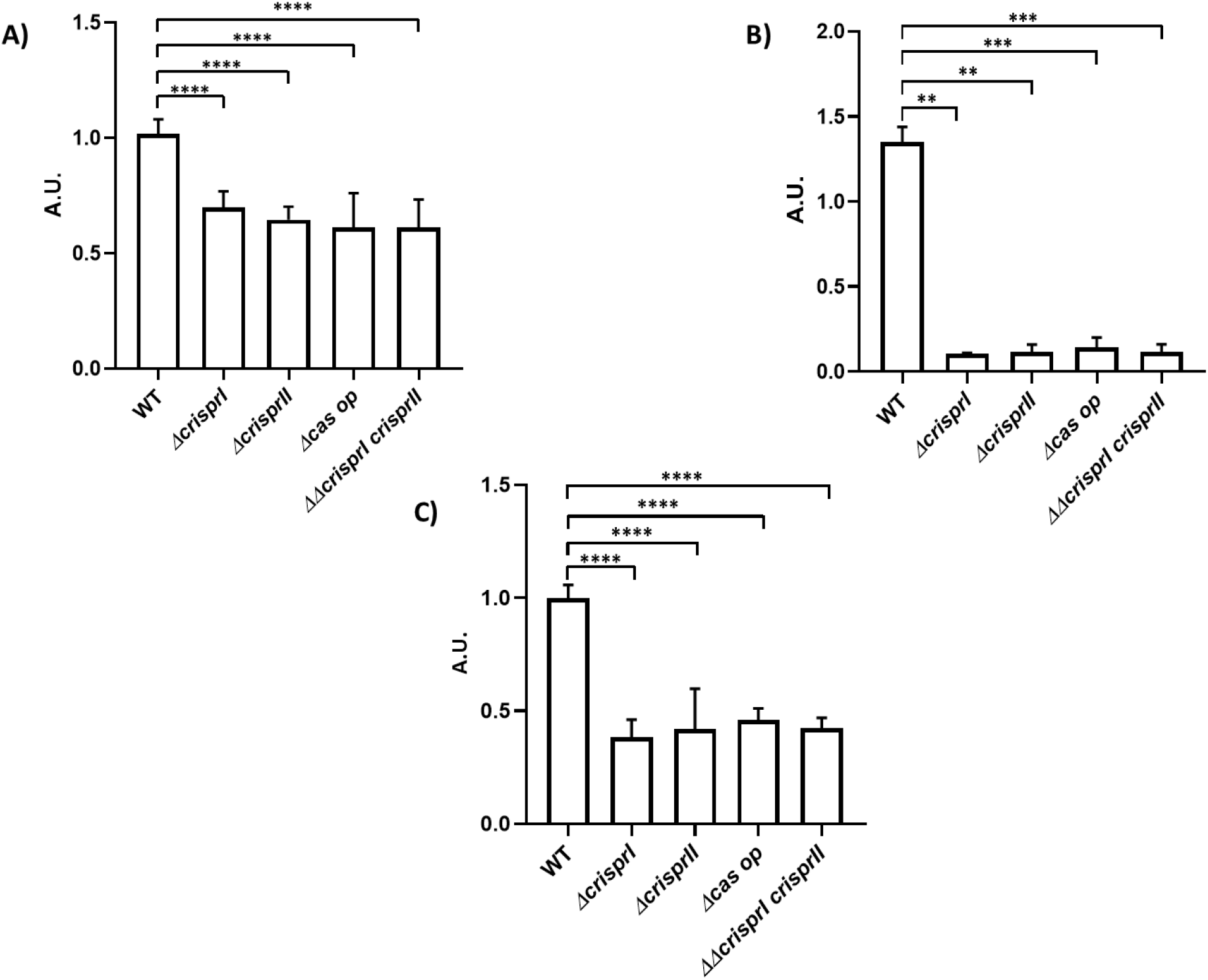
Planktonic culture of the CRISPR-Cas knockout strains showed variations in the productions of key components like curli (A), and cellulose (B). Though thicker than wildtype pellicle biofilm, pellicle biofilms formed by CRISPR-Cas knockout strains were found to be delicate (C). **A.** Production of Curli by the planktonic bacteria of wildtype, CRISPR, and *cas operon* knockout strains was assessed with the help of Congo red depletion assay. The *S.* Typhimurium strain 14028s wildtype (WT), CRISPR (*ΔcrisprI, ΔcrisprII,* and *ΔΔcrisprI crisprII*) and *cas operon (Δcas op*) knockout strains were cultured in LB without NaCl media 48 h, at 25°C, static condition. Congo red depletion was determined by measuring the absorbance of the unbound Congo-red in the supernatant of cultures at 500 nm. The graph represents absorbance at 500 nm for each strain, normalized by absorbance at 500 nm for WT. **B.** The *S.* Typhimurium strain 14028s wildtype (WT), CRISPR (*ΔcrisprI, ΔcrisprII,* and *ΔΔcrisprI crisprII*) and *cas operon (Δcas op*) knockout strains were cultured in LB without NaCl media for 96 h, at 25°C, static condition. Cellulose production in the planktonic culture of wildtype, CRISPR, and *cas operon* knockout strains was quantified by anthrone assay. The graph represents the absorbance of the sample at 620 nm. **C.** The strength of the pellicles were determined by checking the ability of the pellicle to withstand the weight of the glass beads (1 mm, HiMedia). The glass bead weight tolerated by pellicle of each strain was normalized to that of WT. An unpaired t-test was used to determine significant differences between the WT and knockout strains. Error bar indicates SD. Statistical significance: *≤ 0.05, **≤ 0.01, ***≤ 0.001, ****<0.0001, ns = not significant. A.U., arbitrary units.

**Supplementary Figure S9:**
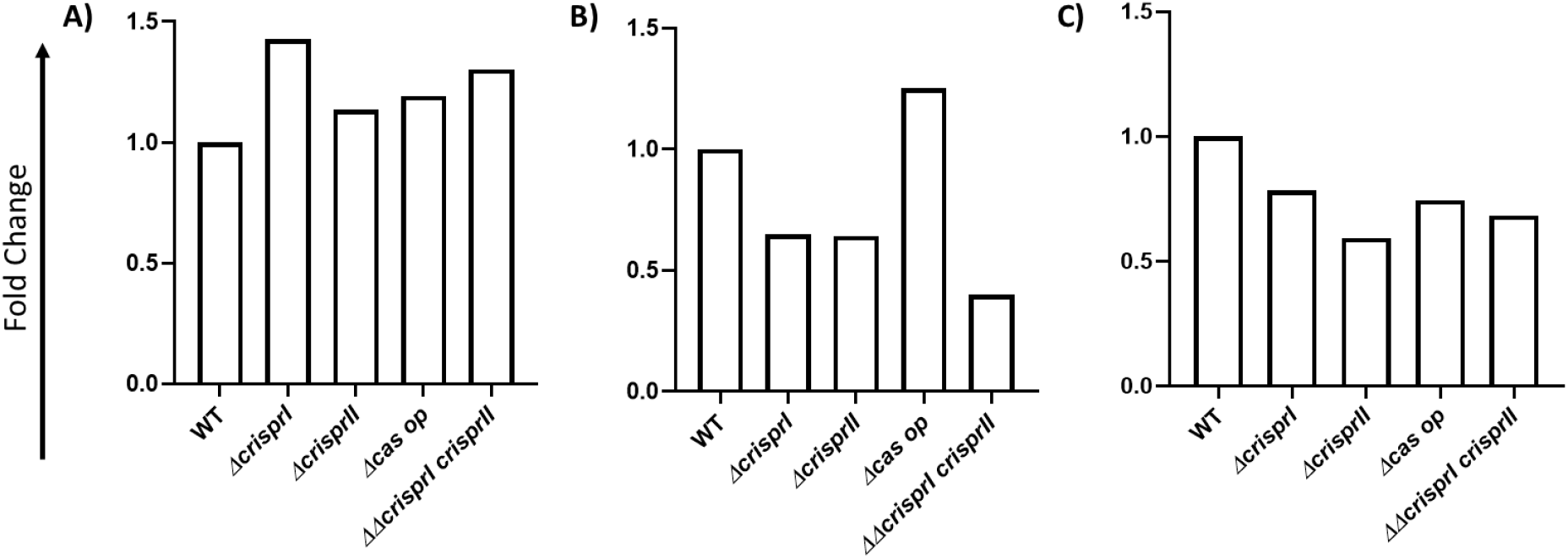
CRISPR-Cas system knockout strains showed differences in the expressions of genes associated with flagellar protein *flgJ* (A), *rfbI* (B), and *bcsA* (C) when compared to WT. The *S.* Typhimurium strain 14028s wildtype (WT), CRISPR (*ΔcrisprI, ΔcrisprII,* and *ΔcrisprI ΔcrisprII*) and *cas operon (Δcas op*) knockout strains were cultured in LB without NaCl media for 24 h, at 25°C, static condition. Total RNA was isolated from bacteria using TRIzol reagent as per the manufacturer's instructions. 1 μg of RNA was used for cDNA synthesis, followed by qRT-PCR. Relative expression of the gene was calculated using the 2^-ΔΔCt^ method and normalized to reference gene *rpoD.*

**Supplementary Figure S10:**
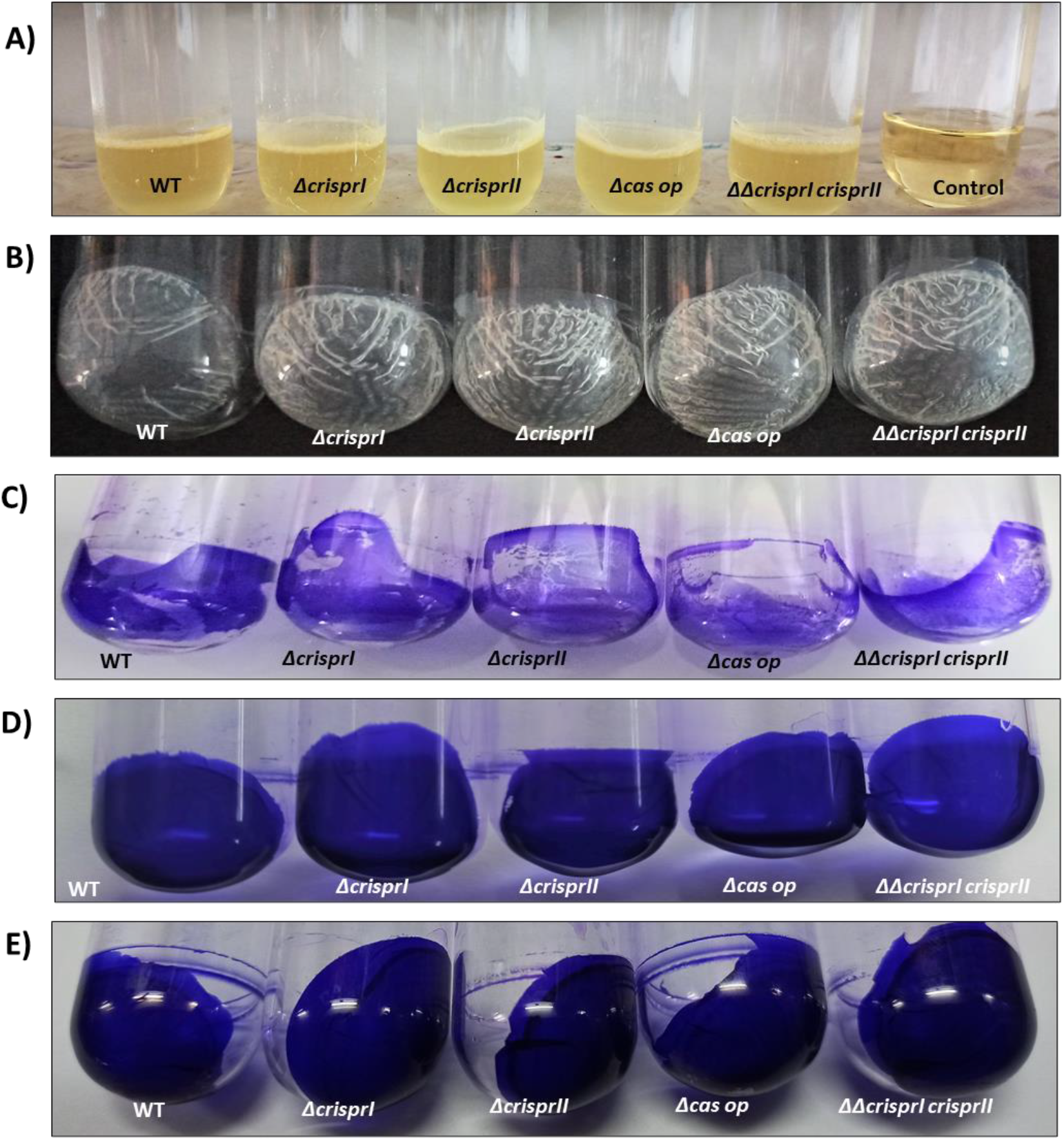
Representative images of Pellicle Biofilms. **A.** Biofilm formation by *S. enterica* subsp. *enterica* serovar Typhimurium 14028s wildtype and CRISPR-Cas system knockout strains at air-liquid interphase (pellicle). **B.** Unstained pellicle biofilm of *S. enterica* subsp. *enterica* serovar Typhimurium 14028s wildtype and CRISPR-Cas system knockout strains. **C.** CV-stained, 24 h pellicle biofilm of *S. enterica* subsp. *enterica* serovar Typhimurium 14028s wildtype and CRISPR-Cas system knockout strains. **D.** CV-stained, 48 h pellicle biofilm of *S. enterica* subsp. *enterica* serovar Typhimurium 14028s wildtype and CRISPR-Cas system knockout strains. **E.** CV-stained, 96 h pellicle biofilm of *S. enterica* subsp. *enterica* serovar Typhimurium 14028s wildtype and CRISPR-Cas system knockout strains.

**Supplementary Figure S11:**
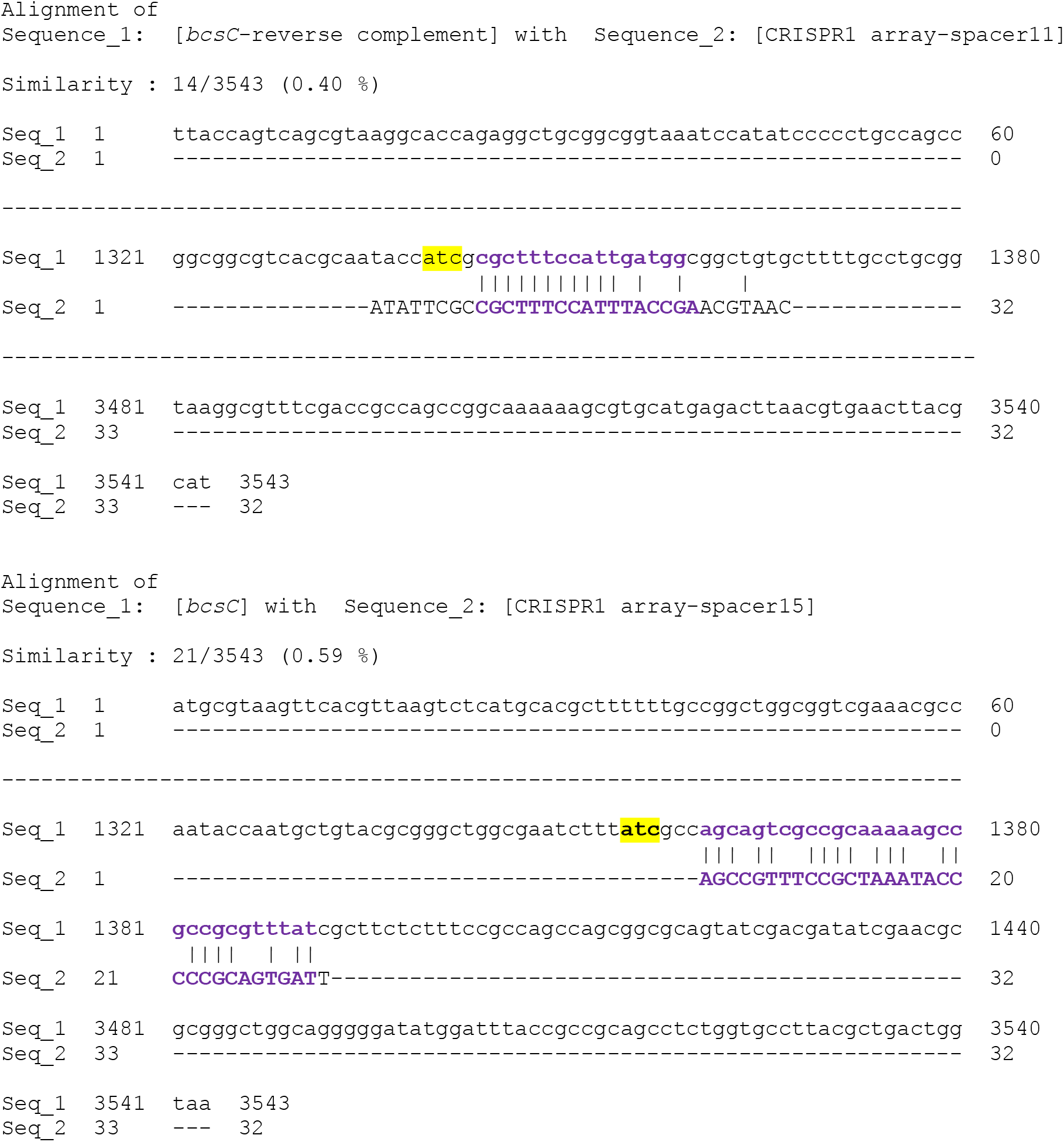
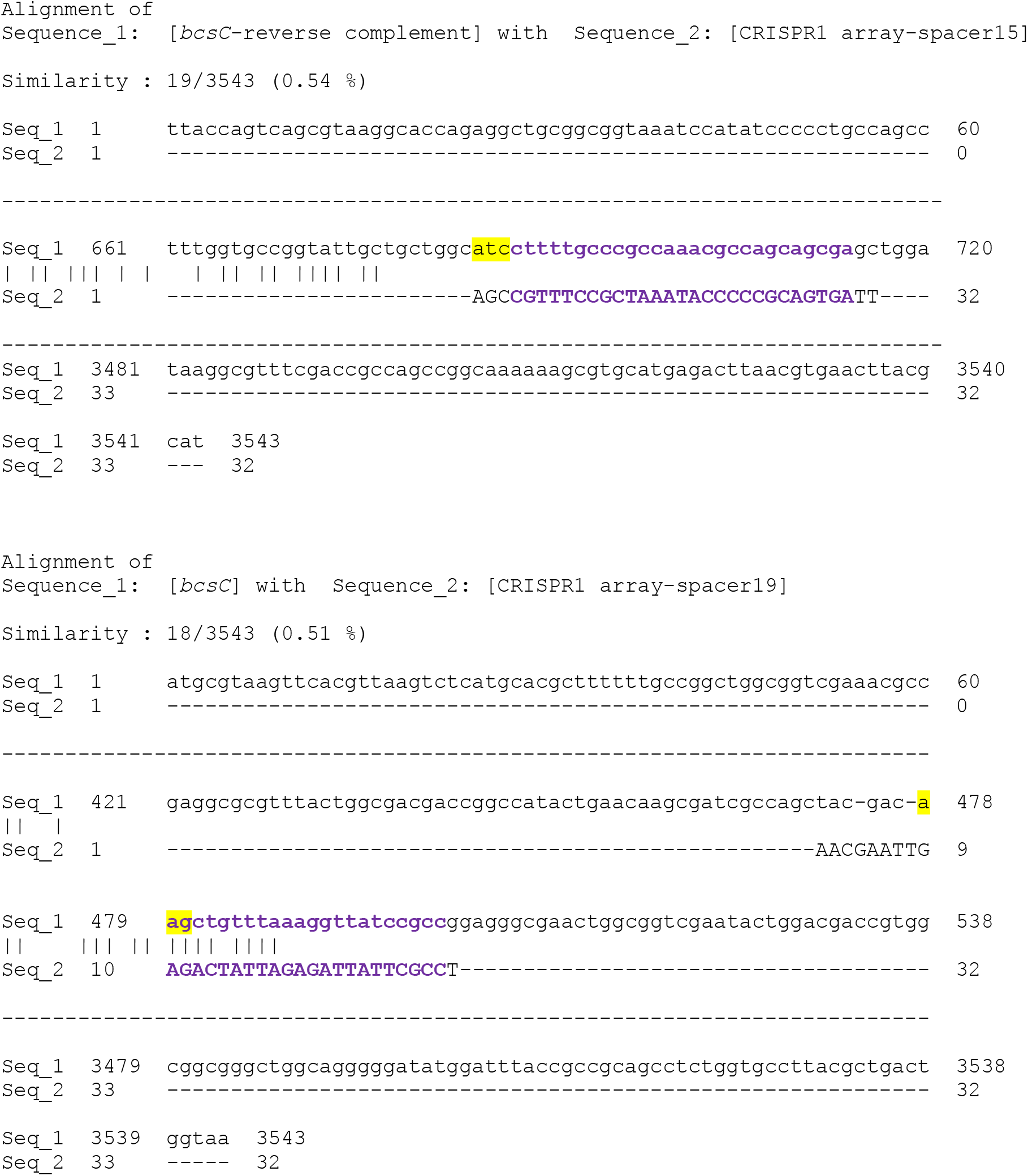
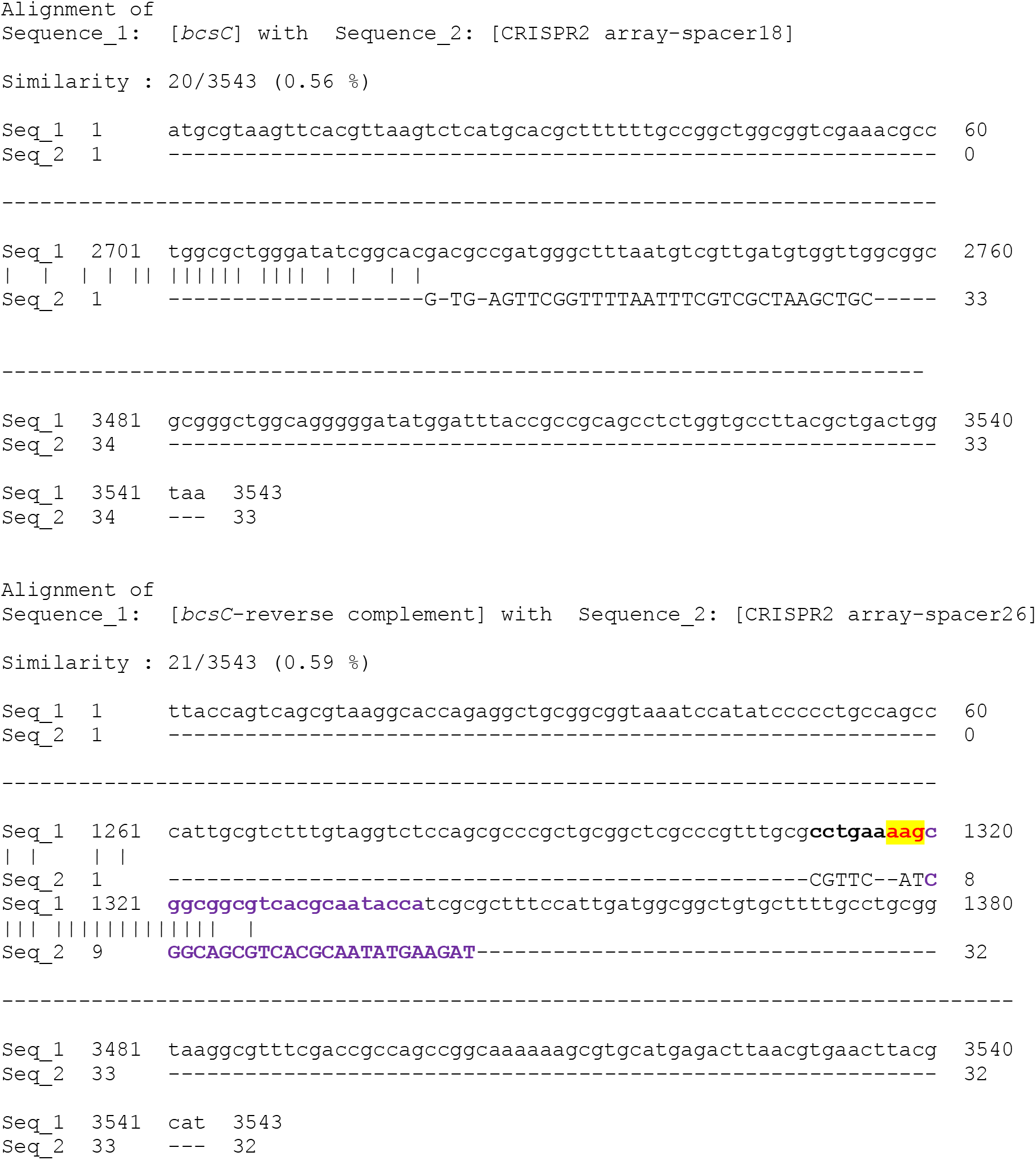
Partial complementarity between spacers (spacer 11, 15 and 19 in CRISPRI array and 18 and 26 in CRISPRII array) and *bcsC* gene. The coding and the reverse complement (template) sequence of the *bcsC* gene were extracted from a complete-genome sequence of Typhimurium str. 14028S, NCBI (GenBank: CP001363.1). The spacer sequences of CRISPRI and CRISPRII arrays were then aligned with coding and reverse complement of *bcsC* gene using serial cloner version 2.6 software. The putative PAM sequences are highlighted in yellow.

